# Fast, Flexible, Feasible: A Transparent Framework for Evaluating eDNA Workflow Trade-offs in Resource-Limited Settings

**DOI:** 10.1101/2025.04.16.649249

**Authors:** Yin Cheong Aden Ip, Elizabeth Andruszkiewicz Allan, Shana Lee Hirsch, Ryan P. Kelly

## Abstract

Environmental DNA (eDNA) analysis enables biodiversity monitoring by detecting organisms from trace genetic material, but high reagent costs, cold-chain logistics, and computational demands limit its broader use, particularly in resource-limited settings. To address these challenges and improve accessibility, we systematically evaluated multiple workflow components, including four DNA extraction methods, two primer sets, three Nanopore basecalling models, and two demultiplexing pipelines. Across 144 workflow combinations tested in an aquarium with 15 fish species, we mapped trade-offs between cost, sensitivity, and processing speed using a hierarchical Bayesian model, to assess where time and resource savings are possible without compromising detection. Workflows using the Qiagen Blood and Tissue (BT) extraction kit provided the highest sensitivity, recovering all 15 species of the sampled fish community within 3–5 hours of Oxford Nanopore sequencing when paired with MiFish-U primer set and high-accuracy (HAC) basecalling. Chelex, an alternative lower-cost extraction method, required extended sequencing (>24 hours) to reach comparable species-detection rates. DirectPCR and QuickExtract offered field-friendly extraction alternatives that achieved comparable recovery in ∼10–12 hours, though their cost-effectiveness varied. While the MarVer1 primer was designed to broaden vertebrate detection, it recovered the same fish species as MiFish-U, though with fewer total reads. Real-time sequencing trials (0–61 hours) revealed that high-efficiency workflows (BT + HAC) reached detection plateaus rapidly, indicating sequencing time can be reduced without sacrificing accuracy. The OBITools4 bioinformatics pipeline enabled automated demultiplexing but discarded more reads than an alternative, ONTbarcoder2.3, which retained low-abundance taxa at the cost of manual curation. Rather than identifying a single “best” workflow, this study provides a transparent decision framework for prioritizing cost, speed, and sensitivity in eDNA applications, supporting scalable, cost-effective eDNA monitoring, in resource-limited settings.

## 1. Introduction

Analysis of environmental DNA (eDNA) is revolutionizing biodiversity monitoring by enabling species detection from trace genetic material in water, air, and sediment, offering a powerful alternative to traditional survey methods (Deiner et al., 2017). While eDNA itself refers to the genetic material shed by organisms into the environment, its utility relies on robust and replicable analytical workflows. By capturing elusive or cryptic taxa that are often overlooked by conventional techniques (Ip et al., 2021), eDNA analysis has expanded the scope of ecological research and conservation efforts. However, its effectiveness depends on key methodological choices, including DNA extraction, primer selection, sequencing accuracy, and bioinformatics processing, which directly influence species detection rates and the comparability of results across studies (Allan et al., 2025 preprint; Mathon et al., 2021; Tsuji et al., 2019).

Beyond these methodological considerations, eDNA work is constrained by logistical and technical challenges, particularly in remote field stations and under-resourced laboratories. Many settings lack access to reagents at a reasonable price, specialized laboratory equipment (e.g., benchtop sequencers, high-speed centrifuges), cold-chain storage, and reliable electricity, limiting the ability to process and analyze eDNA samples effectively (Kranzfelder et al., 2016; Majaneva et al., 2018; Hirsch et al., 2024). Additionally, eDNA metabarcoding workflows require extensive computational processing for analyses including basecalling, demultiplexing, and taxonomic classification. This often exceeds the capacity of field-based or small-scale laboratories (Plewnia et al., 2024). For real-time sequencing platforms, optimizing sequencing duration is equally important, as continued sequencing may reach a plateau where additional runtime provides diminishing gains in species detection (Chang et al., 2024). Primer selection in resource-limited settings presents a critical trade-off: universal primers allow for broader taxonomic detection but may sacrifice species-level resolution depending on the target gene region, while taxon-specific primers can provide enhanced resolution within a group but limit the breadth of taxa detected. Consequently, researchers seeking both broad taxonomic diversity and species-specific data must often sequence multiple marker regions, significantly increasing costs (Ficetola & Taberlet, 2023). Thus, without clear guidance on marker selection, suboptimal primer choice can introduce taxonomic biases and reduce the reliability of biodiversity assessments (Shaffer et al., 2025).

To address these limitations, recent initiatives have focused on democratizing eDNA methodologies, making them more accessible to eDNA-users working with limited budgets and infrastructure. Portable sequencing platforms, such as Oxford Nanopore Technologies’ (ONT) MinION, have been leveraged for on-site sequencing and library preparation (Watsa et al., 2020; Chang et al., 2020; Kirchgeorg et al., 2024), and international standardization efforts are starting to aim to optimize protocols for field-deployable workflows without compromising data quality (Laamanen et al., 2025; Hirsch et al., 2024). Furthermore, a growing suite of user-friendly extraction guidelines (Rieder et al., 2024) has lowered technical barriers and improved accessibility for new practitioners. To maximize the benefit of these advances, here we undertake a systematic evaluation of how methodological choices—from extraction and primer selection to basecalling and demultiplexing—might affect species detection, cost-efficiency, and feasibility.

### DNA Extraction

DNA extraction is one of the most resource-intensive steps in eDNA workflows and influences every subsequent step in the analytical chain. Commercial column-based kits are the most used eDNA extraction methods, with ∼50% of studies relying on Qiagen Blood & Tissue (BT) kits (Tsuji et al., 2019). These kits are favored for their consistency and ease of use; however, their cost and requirement for specialized equipment (e.g. centrifuge) often render them impractical for field applications. In contrast, simplified methods like Chelex-based protocols offer an ultra-low-cost alternative (∼$0.05 per sample; Holman et al., 2019) with minimal reliance on specialized equipment, though their susceptibility to PCR inhibitors can potentially reduce the detection of low-abundance taxa in environmental samples (Walsh et al., 1991; Bracken et al., 2019; Karlsson et al., 2022). Other simplified extraction methods like QuickExtract (QE) and DirectPCR represent intermediate solutions, requiring minimal infrastructure while maintaining moderate efficiency, but their performance can also vary in turbid or inhibitor-rich samples (Majaneva et al., 2018; Lee-Rodriguez et al., 2024; Scriver et al., 2024). Understanding how low-cost, simplified extractions impact species recovery and where optimizations can mitigate losses is key to balancing feasibility of eDNA protocols with accuracy in species detection.

### PCR Primers

Choosing the right primer set is fundamental to eDNA workflows, as it directly impacts detection sensitivity and the taxonomic resolution of generated data. For instance, MiFish-U (Miya et al., 2015) is optimized for fish-specific recovery, while broader-coverage primers like MarVer1 (Valsecchi et al., 2019), target the same 12S region but were designed for wider vertebrate coverage (Doorenspleet et al., 2021; Plewnia et al., 2024; Shaffer et al., 2025).

Notably, MarVer1 is known to amplify human DNA particularly well (Shaffer et al., 2025), potentially complicating analyses in remote or resource-limited settings where strict contamination control is difficult. Different primer designs can influence the balance between taxonomic breadth and target specificity, an important consideration when choosing markers for multi-taxon versus single-group studies.

Beyond taxonomic specificity, primer choice is shaped by target amplicon length, which influences sequencing strategy and analytical feasibility. Both MiFish and MarVer1 amplify fragments in the 150–200 bp range, a commonly targeted length in eDNA metabarcoding due to high PCR efficiency, broad applicability, and widespread use in short-read metabarcoding workflows (Wang et al., 2023; Yang et al., 2024). Finally, while longer amplicons (>650 bp) improve taxonomic resolution and phylogenetic inference, they introduce challenges related to amplification efficiency, library preparation, sequencing depth, and bioinformatics complexities (Chang et al., 2024; Deiner et al., 2017; Kirchgeorg et al., 2024; Tibone et al., 2025). Therefore, primer selection is a critical decision point that must balance specificity, taxonomic breadth, and sequencing constraint to optimize eDNA workflows for different ecological applications.

### Downstream Processing

Post-sequencing processes play a crucial role in eDNA analysis workflows as they influence the accuracy, efficiency, and scalability of taxonomic identification. Wet lab protocol choices, such as indexing strategies, adapter ligation, and barcode designs, can affect demultiplexing efficiency, particularly in ONT workflows where basecalling accuracy affects barcode recognition (Krehenwinkel et al., 2019). Ensuring synergies between sequencing preparation and data processing strategies can help minimize barcode assignment errors and optimize downstream analyses (Petit-Marty et al., 2023).

Unlike Illumina sequencing, which delivers final nucleotide sequences (AGCT) directly, ONT records raw electrical signals (i.e. voltage) that must first be converted into base calls, a process known as basecalling. ONT offers three basecalling models—Fast, High Accuracy (HAC), and Super Accuracy (SUP), which allow users the option to balance computational efficiency and read fidelity. These differences directly impact taxonomic resolution and the likelihood of achieving species-level identification (Wick et al., 2019; Chang et al., 2024). Demultiplexing influences detection outcomes by determining how efficiently with which reads are assigned to their respective samples. Tools such as ONTbarcoder2.3 and OBITools4 handle read errors, indels, and barcode mismatches differently, resulting in trade-offs between automation, read retention, and the need for manual intervention (Chang et al., 2024). While permissive tools may preserve a higher number of reads, they often introduce greater workflow overhead, requiring manual intervention and multiple systems to process many files concurrently, whereas more automated pipelines reduce such labor even if they discard a small proportion of reads. The optimal demultiplexing strategy thus depends on the dataset complexity, available resources, and the desired level of accuracy.

Following demultiplexing, taxonomic classification also influences workflow efficiency, as differences in classification algorithms, reference database completeness, and error tolerance can impact species identification and detection sensitivity. Curated reference databases like MitoFish (Zhu et al., 2023), enhance classification accuracy by providing curated and taxonomically comprehensive resources for species assignment. To support rapid, high-throughput analysis of large ONT datasets, we selected Kraken2 as our taxonomic classifier because of its computational efficiency and compatibility with custom reference databases (Zhu et al., 2023; Bayer et al., 2024; Liu et al., 2024; Lepuschitz et al., 2020; Lu et al., 2022). These bioinformatic decisions are integral to balancing performance and scalability in eDNA metabarcoding workflows, aligning metabarcoding outcomes with the constraints of available computational resources and research priorities.

This study systematically evaluates the trade-offs in eDNA workflows, identifying how methodological “shortcuts” for cost-effectiveness and time-efficiency remain viable and where they introduce meaningful losses in resulting data quality (Figure 1). We compare four DNA extraction methods (Qiagen BT, Chelex, QuickExtract, and DirectPCR) with two 12S primer sets (MiFish-U and MarVer1), three Nanopore basecalling models (Fast, HAC, and SUP), and two demultiplexing pipelines (OBITools4 and ONTbarcoder2.3), resulting in 144 unique workflow combinations tested in a controlled aquarium containing 15 known fish species. In addition to endpoint analyses, we examined species accumulation trends in real-time sequencing by analyzing ONT data at hourly intervals from zero to 61 hours, assessing how sequencing duration can be reduced without sacrificing accuracy. By comprehensively assessing detection accuracy and computational efficiency, we provide a decision framework for researchers to tailor eDNA workflows based on specific priorities, whether they be for maximizing species recovery, minimizing costs, or enabling rapid field-based analyses. Ultimately, these findings contribute to democratizing eDNA-based biodiversity monitoring, ensuring that cost-effective and scalable approaches remain accessible for conservation, invasive species management, and ecological research, particularly in resource-limited settings.

**Figure 1.**
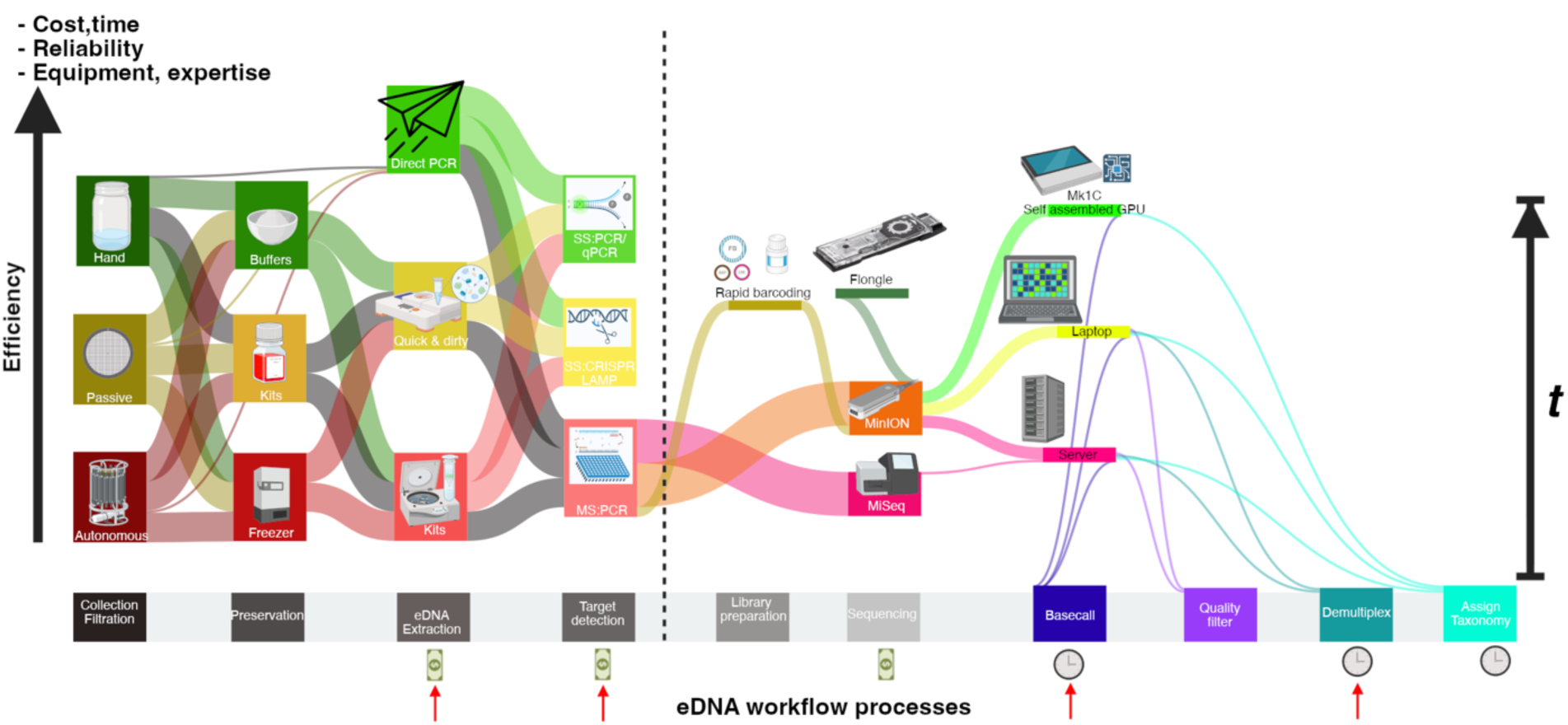
Conceptual overview of eDNA workflows and trade-off considerations. This figure illustrates the eDNA workflow from sample collection to taxonomic assignment, highlighting key trade-offs in cost, time, reliability, and equipment requirements. The vertical axis represents a composite efficiency metric (e.g., cost, time, and resources per taxon detected), with streamlined, field-friendly methods at the top and resource-intensive workflows at the bottom, while the horizontal axis tracks sequential eDNA processing steps. Red arrows indicate the methodological steps tested in this study, i.e. eDNA extraction, target detection (broad vs. specific primers), basecalling models, and demultiplexing workflows; each accompanied by red arrows with cash or clock symbols to denote opportunities for cost savings or time optimization. Sequencing (MinION) and taxonomic assignment (Kraken2) were selected for their affordability and speed but were not directly tested in this study, and thus, do not feature red arrows. These workflow optimizations, including simplified extraction, Nanopore sequencing as a cost-effective alternative to the more common Illumina MiSeq, and accelerated bioinformatics pipelines for taxonomic classification, enhance the feasibility of eDNA analysis, particularly in resource-limited settings.

## 2. Materials and Methods

### 2.1. Experimental Design and Selection of eDNA Extraction Methods

We employed a factorial design to systematically assess the effects of various methodological choices on eDNA metabarcoding outcomes. In total, we processed 12 biological samples (three replicates per extraction across four methods) using two primer sets, three basecalling models, and two demultiplexing pipelines, resulting in 144 unique workflow combinations. This design enabled the evaluation of both individual factor effects and potential interactions. To account for variability, we included biological replicates (three replicate water samples per extraction method) and technical replicates (three PCR replicates per sample, aggregated at the PCR level). This replication strategy facilitated a robust assessment of workflow consistency.

Four DNA extraction methods were evaluated for feasibility and efficiency in resource-limited contexts (Table 1):

1. DNeasy Blood & Tissue (BT) (Qiagen, Hilden, Germany): A widely used column-based approach (Tsuji et al., 2019) that provides high DNA recovery and purity but is expensive and require multiple reagents plus specialized laboratory equipment, such as a high-speed centrifuge.
2. Chelex-100 (Chelex) (Bio-Rad, Hercules, CA, USA): A low-cost resin-based method that can reduce per-sample costs to approximately $0.05 (Bracken et al. 2019) and require minimal equipment but sometimes exhibiting reduced sensitivity to rare taxa due to inhibitor presence (Karlsson et al. 2022).
3. QuickExtract (LGC Biosearch Technologies, Middleton, WI, USA): An enzymatic lysis approach requiring minimal equipment and moderate cold storage. It has proven effective for small insect exuviae (Kranzfelder et al. 2016) and greenhouse arthropod eDNA (Lee-Rodriguez et al. 2024), but occasionally retaining inhibitors from turbid samples (Majaneva et al. 2018).
4. DirectPCR/Centrifugal Dialysis (MilliporeSigma, Burlington, MA, USA): Relies on concentrating resuspended water into smaller volumes and enabling direct amplification of DNA from environmental samples without purification. It does not require cold storage and is promising for field applications, though performance can vary in low-DNA contexts, and it does need a mini-centrifuge (Ip et al., 2024; Kirtane & Deiner 2024).

**Table 1.** Comparison of cost, processing time, and equipment requirements for each extraction method. Note: “Time” is expressed in relative units representing processing speed.

| Extraction Method | Cost | Time | Reagent Requirements | Cold Storage | Specialized Equipment | Yield | Purity |
| --- | --- | --- | --- | --- | --- | --- | --- |
| BT | \$\$\$\$ | 4 | High | No | High-speed centrifuge | High | High |
| Chelex | \$ | 3 | Low | No | Heat block | Low | Low |
| QuickExtract | \$\$\$ | 2 | Low | Moderate | Heat block | Medium | Low |
| DirectPCR | \$\$ | 1 | None | None | Mini-centrifuge | Medium | Low |

### 2.2. Sample Collection, Filtration, and Preservation

To provide a controlled setting for evaluating different eDNA methods and species detection rates, water samples were collected from the Seattle Aquarium (Seattle, WA, USA). The facility houses a 450,000 L mixed-species aquarium containing 19 known fish species (see Supplementary Table S1), which provides a standardized setting for workflow comparisons.

A 20 L water sample was taken from the surface of the aquarium tank using a 30 L carboy pre-treated with 10% bleach, rinsed with DI water, and handled with gloves to minimize contamination. Filtration was performed on site using a Smith-Root Citizen Science Sampler and filtration housings equipped with 5.0 µm mixed cellulose ester (MCE) filters (Allan et al., 2023). Additionally, five field-negative filtration blanks (1 L of Milli-Q water each) were processed using the same system. In total, 15 L of aquarium water was filtered (1 L per replicate) within 30 minutes to ensure all samples were filtered and processed under similar conditions, minimizing potential biases from DNA degradation. To maintain homogeneity, the 30 L carboy was thoroughly shaken before each transfer. Filters were then stored on ice for transport to the laboratory for immediate DNA extraction.

### 2.3. Sample Processing, eDNA extraction and Direct PCR Preparation

Before DNA extraction, filters were carefully transferred with sterile, bleach-treated forceps into 5 mL tubes containing 1.5 mL of Milli-Q water. The filters were randomized across DNA extraction treatments to include three biological replicates of aquarium water and one field blank per extraction method. Each filter was vortexed for 1 minute to resuspend eDNA. For DirectPCR, 500 µL of the resuspended water was transferred to a 50 kDa molecular-weight-cutoff Amicon spin column and centrifuged at 14,000 g for 1 minute each, repeated three times to process the total volume of 1.5 mL. The resulting concentrate (50–100 µL) was eluted by inverting the spin column and centrifuging it at 1000 g for 3 minutes, then used directly for PCR. In contrast, for BT, Chelex, and QuickExtract, manufacturer protocols were followed with minor eDNA-specific optimizations (see Supplementary Material 1). Extractions were performed in two sequential batches: BT and DirectPCR were processed simultaneously, followed by QuickExtract and Chelex. All extractions were completed within 2–4 hours collection. Extracts were stored at −20°C until PCR amplification and sequencing, which occurred within three months.

### 2.4. Library Preparation and Sequencing

Amplicons were generated using two primer sets, MiFish-U (a 12S primer targeting fish; Miya et al., 2015) and MarVer1 (a broader 12S primer for vertebrates and metazoans; Valsecchi et al., 2020). The primer sequences were as follows: MiFish-U Forward: 5’-GCCGGTAAAACTCGTGCCAGC −3’; MiFish-U Reverse: 5’-CATAGTGGGGTATCTAATCCCAGTTTG −3’; MarVer1 Forward: 5’-CGTGCCAGCCACCGCG −3’; MarVer1 Reverse: 5’-GGGTATCTAATCCYAGTTTG −3’. To differentiate samples and reduce sample multiplexing errors, 96 unique molecular identifiers (UMIs – elsewhere called indices or tags; see O’Donnell et al. 2016) were designed and attached to the 5’ end of each primer set. These UMIs were generated with a custom Python script that enforced sequence constraints to enhance error resilience in Nanopore sequencing (Supplementary Material 2). Each UMI was structured as a 14 bp sequence, with a fixed “CC” prefix and suffix, with an internal 10-bp region composed of A, T, and G bases (Eric Coissac, pers. comm.). To ensure robust sample differentiation and minimize misassignments, we enforced a minimum Hamming distance of 3 bp between UMIs, ensuring at least three substitutions difference, thus, minimizing the risk of incorrect read assignments due to single-base substitutions. Additionally, we applied a minimum Levenshtein distance of 3 bp, which accounts for insertions and deletions, further increasing UMI integrity in nanopore sequencing. This UMI design enhances sequencing accuracy, allowing reliable sample identification even in the presence of sequencing errors.

Each PCR reaction was performed in triplicate with a total volume of 20 µL, consisting of 10 µL 2× Phusion Mix (Thermo Fisher Scientific), 0.6 µL DMSO (3% final concentration), 0.5 µL BSA (0.5 µg/µL final concentration), 3.9 µL molecular-grade water, 0.5 µL each of forward and reverse primers (10 µM), and 5 µL of eDNA extract/DirectPCR concentrate. PCR cycling conditions followed Shaffer et al. (2025). Amplicons were pooled and purified using AMPure XP beads (Beckman Coulter) at a 0.7× ratio to remove smaller non-target fragments. Library preparation was carried out following the SQK-LSK-114 ligation sequencing kit protocol (Oxford Nanopore Technologies). The completed library was loaded onto an R10.4.1 MinION flow cell and sequenced on a MinION MK1B platform using MinKNOW v24.06.16 (Windows 11), with the run lasting up to 61 hours. In our study, ONT sequencing produced reads that closely match the amplicon length (approximately 150–200 bp), ensuring that our field-deployable approach remains consistent with traditional short-amplicon methods

### 2.5. Basecalling Models

Raw POD5 reads were basecalled using Dorado v0.9.1 to evaluate three Nanopore basecalling modes: Fast, High-Accuracy (HAC), and Super Accurate (SUP). Basecalling was performed on two hardware setups: (i) An Apple MacBook Pro (2023) with an M3 Max Chip, to assess accessibility and processing times on a more consumer-friendly, widely available system; (ii) An MSI Raider 18 HX Gaming Laptop with an NVIDIA GeForce RTX 4090 GPU, which was used for all final analyses due to its higher computational power. Each dataset was basecalled using all three models for comparison. While only run times were recorded for the MacBook M3, this comparison provides insight into the performance differences between Apple silicon processors and high-end NVIDIA GPUs, helping users weigh trade-offs between hardware accessibility and computational efficiency.

### 2.6. Demultiplexing Pipelines

Demultiplexing was performed using two distinct tools designed for ONT sequencing: ONTBarcoder2.3 (Srivathsan et al., 2024), a commonly used program to process amplicon barcoding libraries on Nanopore platforms, and OBITools4 (https://obitools4.metabarcoding.org), which was tested here as an alternative indel-aware tool.

ONTBarcoder2.3 is widely adopted for its ability to handle barcode-based demultiplexing of Nanopore amplicon data, including detection of self-ligated reads which occur when multiple amplicons become joined during sequencing but are initially processed as a single read. These reads are recognized and split, allowing more usable sequences to be retained for downstream analysis (Chang et al., 2024). In our workflow, ONTBarcoder2.3 was run with a minimum read length threshold of 200 bp and an expected barcode length of 250 bp, while other parameters were kept at default (Supplementary Material 3). The standard genetic code was selected, and only sequences deviating by up to 2 bp from the tag sequence were retained. This threshold was supported by the tag design, which ensured a minimum difference of ≥3 bp between barcode tags to reduce read misassignments. However, ONTBarcoder2.3 is graphical user interface (GUI) based and does not support command line execution, limiting its utility in high-throughput workflows.

OBITools4 was evaluated in this study for its ability to handle insertions and deletions (indels) inherent to Nanopore sequencing data. For OBITools4, we used the *obimultiplex* module, specifying an indel-aware demultiplexing process with a maximum mismatch threshold of two base pairs (Supplementary Material 4). The output from OBITools4 is a single concatenated FASTQ file, with sample information embedded in each sequence header. To organize the data at the sample level, we implemented a custom Python script to parse the sequence headers and split the reads into separate FASTQ files corresponding to individual samples.

To ensure accurate separation of reads originating from each primer set, demultiplexing was performed twice, once for MiFish-U and once for MarVer1. Both programs produced outputs that had been trimmed of primers and UMIs as part of the demultiplexing process, ensuring that only high confidence reads progressed downstream for taxonomic classification.

### 2.7. Taxonomic Classification

To assign species annotations, all sequence classification was conducted with Kraken2 v2.1.2 using a custom MitoFish database (Wood et al., 2019; Zhu et al., 2023). Kraken2 employs a k-mer–based approach that is computationally more efficient than BLAST (Lepuschitz et al., 2020; Lu et al., 2022), thus making it more practical for field deployment and for users with limited computational resources (Plewnia et al., 2024). Recent simulation studies further suggest that Kraken2, when paired with a comprehensive or curated database and a modest confidence threshold (0.05–0.2), achieves a balance between sensitivity and specificity (Liu et al., 2024; Bayer et al., 2024).

For this study, a custom Kraken2 database was constructed by downloading curated mitochondrial sequences for fish species, specifically targeting 12S and COI gene regions, from the MitoFish repository (http://mitofish.aori.u-tokyo.ac.jp). Each sequence was assigned a valid NCBI taxonomic identifier and integrated with the NCBI taxonomy database using the *kraken2-build* function. The database was validated using *kraken2-inspect* and cross-verified against BLAST-based classifications to ensure accuracy.

To classify sequences, a confidence score threshold of 0.1 was applied, and only reads mapped to Actinopterygii in both the MiFish-U and MarVer1 datasets were retained for downstream analysis. Raw read compositions and relative abundances from the Kraken2 results were used for subsequent statistical analyses. Potential genus-level ambiguities, such as those observed in *Sebastes* spp., were manually flagged for further verification (Supplementary Table S1). Given that the target species list was predefined based on the aquarium species community, the absence of some reference sequences in external databases was unlikely to impact species identification significantly.

### 2.8. Real-Time Sequencing

In addition to the consolidated POD5 files mentioned in Section 2.5, the hourly POD5 data from the 61-hour run were processed separately to enable a time-resolved analysis of eDNA detection. After each hourly interval, the corresponding POD5 file was transferred to the MSI Raider 18 HX laptop for basecalling (Fast, HAC, SUP), followed by OBITools4 demultiplexing (leveraging its command-line automation) and Kraken2 classification as described in Section 2.7. This produced 61 hourly snapshots of eDNA readouts, enabling the construction of species accumulation curves that identified the minimum run time needed to recover all known aquarium species. Additionally, species detection frequencies were tracked hourly to assess how abundance and sequencing duration influenced detection frequencies.

### 2.9. Statistical Analysis and Visualization

All statistical analyses were conducted in R Studio (Version 2024.04.1+748, R version 4.2.0) to evaluate how extraction method, primer choice, basecalling model, demultiplexing strategy, and sequencing duration influenced eDNA-based species detection. Raw DNA read counts for the known 15 aquarium species, obtained from Kraken2 outputs, were log₁₀-transformed to reduce skewness and used for bar plots or summary metrics.

To further evaluate how each workflow component influences species detection and compositional proportions, we applied a hierarchical Bayesian model using the zero-and-one inflated Dirichlet (ZOID) framework (Jensen et al., 2022), which allowed us to carry out a straightforward regression analysis on compositional data. Specifically, we employed a partial-aggregation approach by averaging PCR replicates but keeping biological replicates (bio_rep) as distinct independent observations, to estimate factor-level effects on species detection probabilities. Within each sample, proportions were derived by dividing each species’ read count by the total reads of all 15 target species, ensuring the sum of proportions equaled one. In the ZOID design matrix, Qiagen BT extraction, MiFish-U primers, HAC basecalling, and ONTbarcoder2.3 demultiplexing were set as the baseline reference levels. In addition, we compared this partial-aggregation with a fully aggregated approach, in which all replicates (biological and PCR) are averaged post-normalization. This comparison allowed us to assess the impact of aggregation strategy on model outputs; the fully aggregated analysis is presented in Supplementary Material 5, while the partial-aggregation approach is used as our primary analysis because it better preserves inherent biological variability. We generated bivariate, forest, and box plots to visualize factor-level effects across species detections, extracted posterior coefficients for each species-factor combination, and performed posterior predictive checks to model fit and zero-inflation mismatches. To capture potential nonlinear trends in species accumulation over sequencing time, we fitted Generalized Additive Models using the mgcv package (Wood, 2001) with sequencing time as a thin-plate regression spline smooth term.

Hierarchical Bayesian inference was carried out with the zoid package (Jensen et al., 2022), which runs Stan under the hood via rstan (McElreath, 2018). Nonlinear species-accumulation trends were fitted with GAMs using mgcv (Wood, 2001). Data wrangling and visualization relied on the tidyverse suite (Wickham et al., 2019), including dplyr, tidyr, purrr, and stringr. We used ggplot2 and the ggsci extension for plotting, and ggpubr to add statistical annotations.

## 3. Results

Across all workflows, 17 of the 19 expected fish species were detected. Two species, *Rhinogobiops nicholsii* and *Sebastes nebulosus*, were not detected in any workflow, likely due to low eDNA concentrations. BLAST analyses indicate that both primers align well with the 12S region in *Sebastes nebulosus*, which is present in the MitoFish database, suggesting that primer mismatches are unlikely to explain its non-detection. In contrast, *Rhinogobiops nicholsii* is absent from both MitoFish and GenBank databases, precluding in-silico verification of primer compatibility; its non-detection is likely attributed to low template levels. Additionally, two species pairs were grouped due to near-identical 12S sequences for both primer sets, limiting taxonomic resolution: (i) *Hemilepidotus hemilepidotus* with *Hemilepidotus spinosus*, and (ii) *Sebastes flavidus* with *Sebastes diaconus*. Notably, both *Hemilepidotus spinosus* and *Sebastes diaconus* are also missing from the MitoFish database, which may have contributed to the merging of these taxa. This resulted in a final curated dataset of 15 uniquely identifiable species, or Operational Taxonomic Units (OTUs) (see Supplementary Table S1). We consider this count as our maximum number of species reasonably detectable under the constraints of the primer choice, reference database completeness, and the resolution limits of the 12S marker.

The most effective workflow combined DNA extraction with Qiagen BT, MiFish-U primers, HAC/SUP basecalling, and demultiplexing by ONTBarcoder2.3. This combination consistently recovered all 15 OTUs and achieved full detection within 3–5 hours of sequencing (Figure 4,5). However, DirectPCR and QuickExtract also performed reliably in many conditions, demonstrating mid-tier read yields that can be advantageous in low-resource or high-throughput contexts. These results provide a performance benchmark for comparing methodological trade-offs in the sections that follow.

### 3.1. Read recovery and species detection

#### 3.1.1. Effect of extraction method on read depth and species detection

Among the extraction methods, Qiagen BT consistently yielded the highest read counts across all conditions (Figure 2), with BT-extracted samples producing ∼420,000 raw reads per sample replicate in the best workflows. The highest recovery was observed under the SUP basecalling + MiFish-U primers + ONTbarcoder2.3 workflow. In contrast, Chelex consistently exhibited the lowest read counts, with most samples receiving fewer than ∼1,000 reads, reducing the ability to recover low-abundance taxa across all workflows. The best-performing Chelex workflow (HAC + MiFish-U + ONTbarcoder2.3) recovered only ∼3,000 reads per run, nearly an order of magnitude lower than the lowest-throughput Qiagen BT workflows. DirectPCR and QuickExtract yielded intermediate read depths, typically ∼10,000–100,000 reads, offering moderate yields that occasionally approached BT levels when paired with optimal primer and basecalling choices.

**Figure 2.**
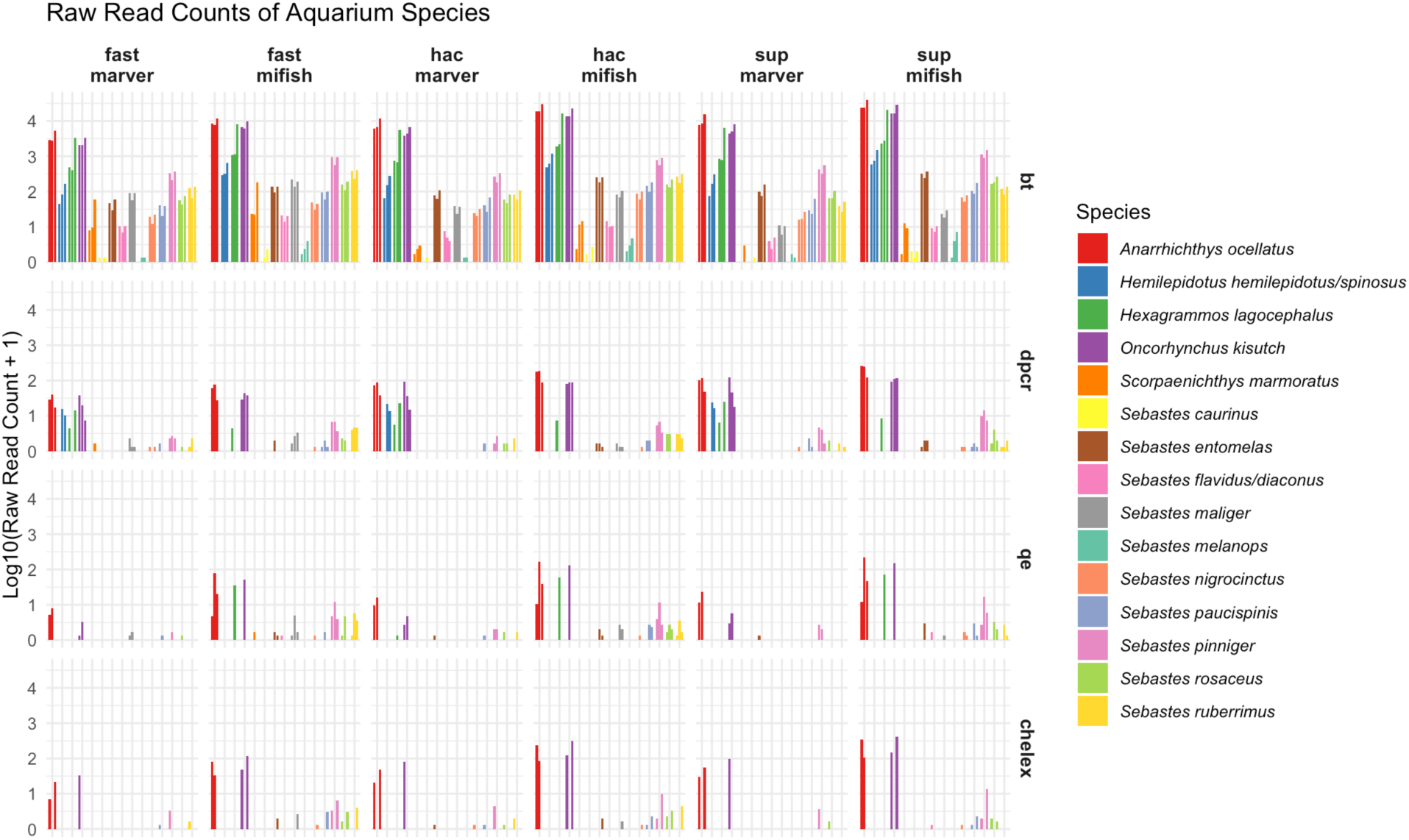
Read Counts Across Workflow Variables in eDNA Analysis. Bar plots display the log_10_-transformed raw read counts of aquarium species detected across various workflow components, including basecalling models (FAST, HAC, and SUP), primer sets (MiFish-U and MarVer1), and DNA extraction methods (BT, Chelex, QuickExtract, and DirectPCR). Only results from the ONTbarcoder2.3 demultiplexing approach are shown here; full results comparing OBITools4 are provided in the supplementary materials. Biological replicates (bt1, bt2, bt3) are shown separately while PCR replicates (a, b, c) are averaged within each biological replicate. Each species appears as up to three side-by-side bars, or fewer if not detected in all replicates).

These patterns are further illustrated in the bivariate comparison of predicted detection proportions from ZOID coefficients (Figure 3, Table 2), where extraction methods show distinct relationships to the baseline BT workflow. Chelex samples consistently fall below the diagonal identity line, demonstrating uniformly reduced detection across all species — a pattern suggesting systematic DNA loss or inhibitor carryover that affects all taxa similarly regardless of abundance. In contrast, DirectPCR samples cluster tightly around the diagonal, indicating performance comparable to BT for most species with only minor variations. QuickExtract shows an intermediate pattern, indicating reduced detection efficiency for certain taxa relative to BT (e.g., *Sebastes maliger*). Notably, Chelex results changes when biological replicate variability is preserved, highlighting that it is highly variable and unreliable compared to BT. DirectPCR and QuickExtract showed neutral to mildly positive coefficients across several species, underscoring their potential as practical, cost-effective alternatives when maximum sensitivity is not critical.

**Figure 3.**
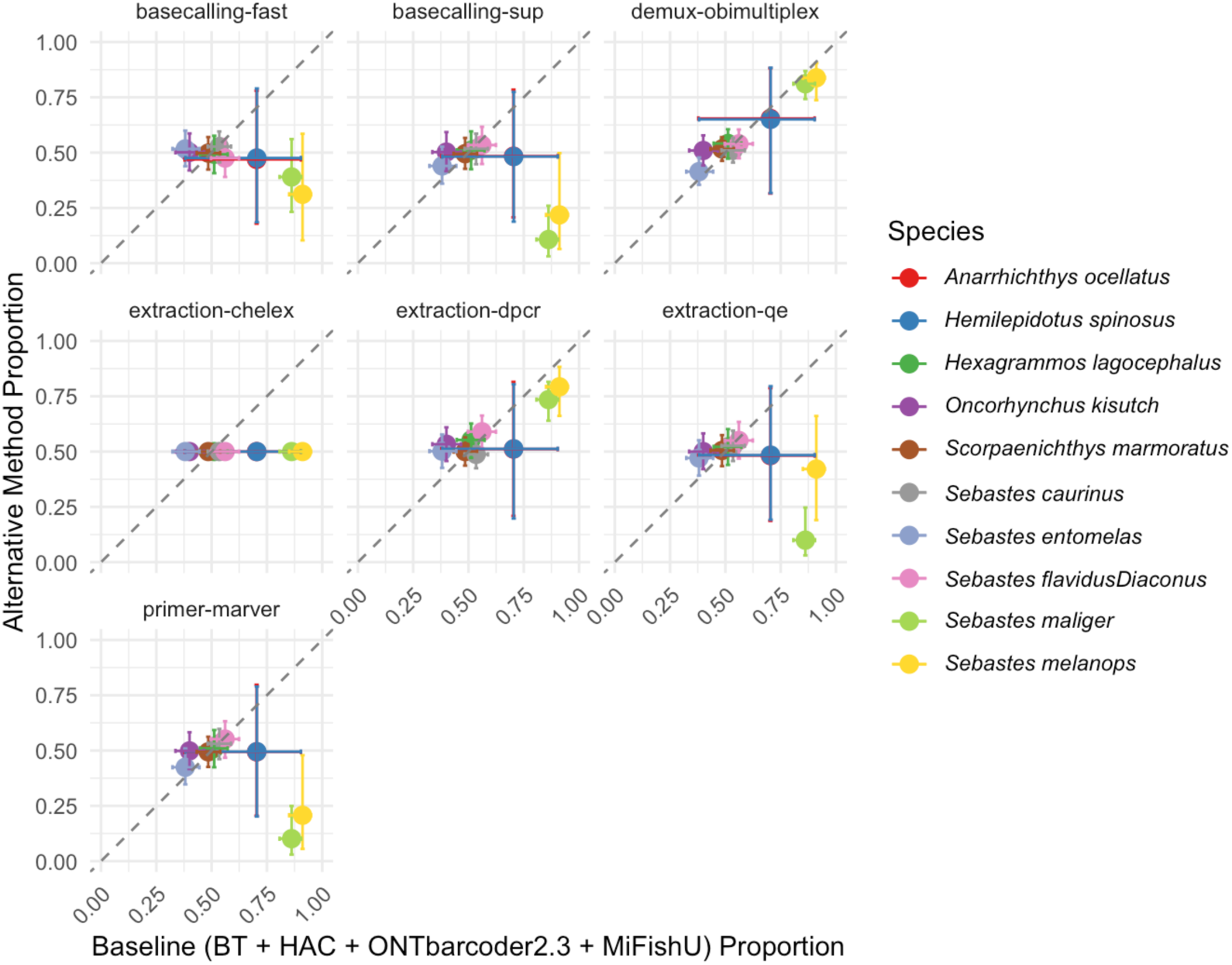
Bivariate Comparison of Predicted Species Detection Proportions. Each panel shows the predicted detection proportion for an alternative method (y-axis) compared to the baseline configuration (BT + HAC + ONTbarcoder2.3 + MiFishU; x-axis). Points represent the posterior mean predicted proportion for each species, and error bars represent the 95% credible intervals. The dashed diagonal line (y = x) indicates equal detection between the baseline and alternative methods. Points above the diagonal denote higher detection probability under the alternative method, while points below indicate lower detection probability.

**Table 2.** Example output of Zero-and-One Inflated Dirichlet (ZOID) posterior estimates for a single species (*Anarrhichthys ocellatus*), showing the posterior mean and 95% credible intervals for methodological factors relative to the baseline workflow (Qiagen BT, MiFish-U, HAC, and ONTbarcoder2.3). Full results for all 15 species and all factor combinations are provided in Supplementary Material 5.

| Species | Factor | Posterior Mean | 95% Credible Interval (Lower, Upper) |
| --- | --- | --- | --- |
| *Anarrhichthys ocellatus* | (Intercept) | 0.8637 | (-0.5009, 2.2376) |
|  | bio_rep2 | -0.0891 | (-1.5387, 1.3084) |
|  | bio_rep3 | 0.2489 | (-1.1689, 1.7096) |
|  | demux-obimultiplex | 0.6384 | (-0.7768, 2.0065) |
|  | primer-marver | -0.0241 | (-1.3516, 1.3701) |
|  | basecalling-fast | -0.1230 | (-1.5233, 1.2630) |
|  | basecalling-sup | -0.0648 | (-1.3421, 1.2925) |
|  | extraction-dpcr | 0.0453 | (-1.3289, 1.4842) |
|  | extraction-qe | -0.0731 | (-1.4720, 1.3047) |
|  | extraction-chelex | 0 | (0, 0) |

#### 3.1.2. Effect of primer, basecalling algorithm, and demultiplexing on read depth and species detection

Primer selection strongly influenced read recovery, with MiFish-U consistently outperforming MarVer1 across workflows (Figure 2). MiFish-U samples generated significantly higher raw read counts, reaching ∼50,000 to 300,000 reads per sample replicate compared to ∼3,000–20,000 from MarVer1 (Figure 3). Notably, *Sebastes rosaceus* and *Sebastes ruberriumus* showed particularly reduced detection with MarVer1 primers. Both primer sets recovered 15 fish species in the aquarium, but MiFish-U demonstrated greater sensitivity for fish detection, which aligns with expectations as MarVer1 amplifies a broader range of vertebrates (Shaffer et al., 2025), diluting fish read counts.

Basecalling algorithms showed performance stratification in both read depth (Figure 2) and species detection (Figure 3). Fast basecalling prioritized speed but exhibited notably lower sensitivity for rare taxa, HAC balanced speed and accuracy, while SUP achieved the highest read quality (mean Q-score 15.4) and read retention, with many samples exceeding ∼100,000 raw reads per replicate (Table 3). Species detections were remarkably similar between HAC and SUP models, indicating that the extra computation time required for SUP does not necessarily translate into higher detection rates. Thus, HAC offers an optimal balance between computational efficiency and detection sensitivity for most applications.

**Table 3.**
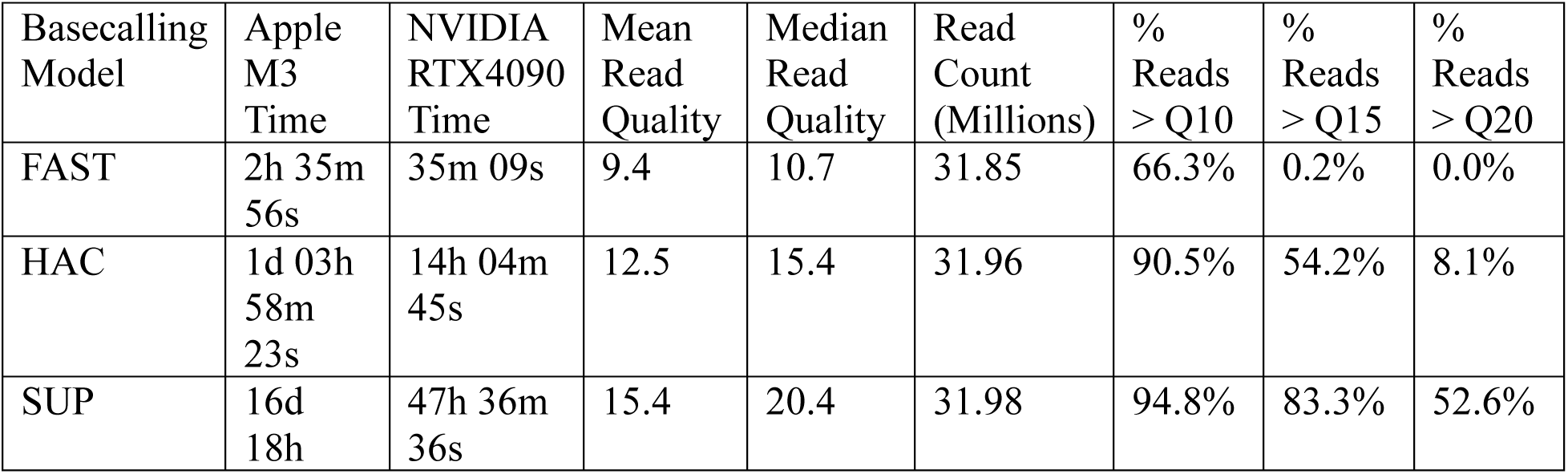
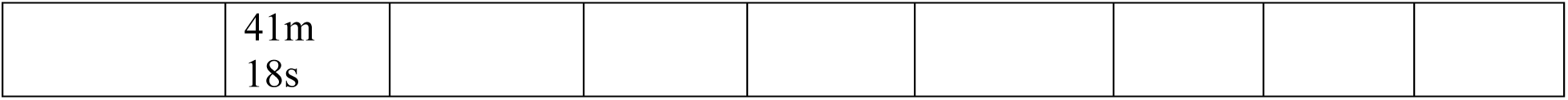
Comparison of Basecalling Models for Read Quality and Retention.

The choice of demultiplexing tool also influenced read retention and species detection (Supplementary Figure S1). In the bivariate comparison plots (Figure 3), OBITools4 and ONTbarcoder2.3 (baseline) show generally similar detection proportions across most species. However, ONTbarcoder2.3 retained more reads overall, improving the detection of rare taxa, particularly in workflows using Chelex-extracted samples or FAST basecalling. These subtle differences in read retention may affect detections for certain species (Supplementary Figure S5).

Overall, BT extraction, MiFish-U primers, and ONTbarcoder2.3 demultiplexing with HAC/SUP basecalling yield the highest detection odds across most taxa (Figure 2, 3). Chelex extraction or MarVer1 primers showed reduced performance for most species, while other factor levels exhibited milder effects. DirectPCR and QuickExtract, though not matching BT in all scenarios, still demonstrated reasonable performance when paired with MiFish-U and either HAC or SUP, underscoring their potential as cost-effective alternatives for many applications (Table 4).

**Table 4.**
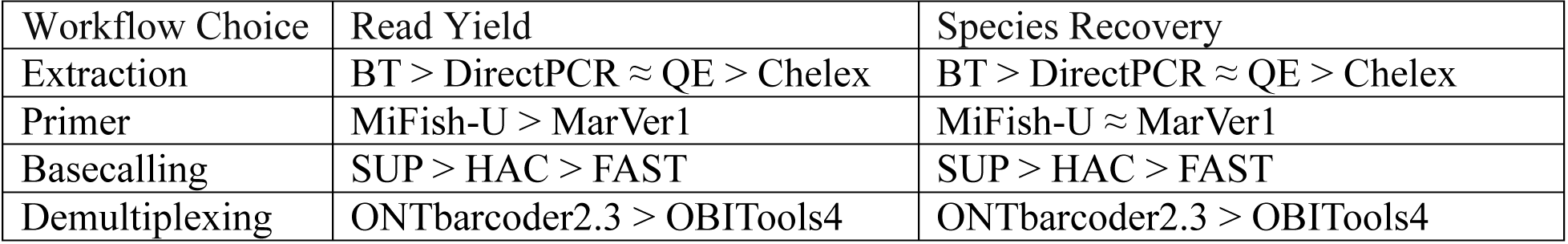
Workflow Performance Summary Comparison of extraction, primers, basecalling, and demultiplexing based on read yield and species recovery.

| Workflow Choice | Read Yield | Species Recovery |
| --- | --- | --- |
| Extraction | BT > DirectPCR ≈ QE > Chelex | BT > DirectPCR ≈ QE > Chelex |
| Primer | MiFish-U > MarVer1 | MiFish-U ≈ MarVer1 |
| Basecalling | SUP > HAC > FAST | SUP > HAC > FAST |
| Demultiplexing | ONTbarcode2.3 > OBITools4 | ONTbarcode2.3 > OBITools4 |

### 3.2 Time, computational power, and other considerations for workflows

#### 3.2.1 Basecalling algorithm speed and computation time

FAST basecalling prioritized speed, processing a full dataset in ∼30 minutes on an NVIDIA RTX 4090 and ∼2 hours on an Apple M3 Max (Table 3). HAC provided a balance between speed and accuracy, requiring ∼14 hours on an NVIDIA RTX 4090 and over a day on the Apple M3 Max. SUP achieved the highest read accuracy; however, this improvement came at a significant computational cost, requiring ∼50 hours on an RTX 4090 and over two weeks on an Apple M3 Max. The number of reads recovered and read quality scores based on the basecalling algorithm affected species detections, though the jump from HAC to SUP did not seem to be worth the nearly 4x computation time in the NVIDIA RTX 4090 or the 14x computation time on the Apple M3 Max, given only marginal gains in species detections.

#### 3.2.2 Real-Time Species Accumulation Over 61 Hours

Cumulative species accumulation curves illustrate the rate at which the 15 unique species OTUs were detected over the course of a 61-hour sequencing run (Figure 4, 5). Sequencing duration strongly influenced detection probabilities, particularly for rare taxa such as *Sebastes ruberrimus*, which required extended sequencing times to reach stable detection thresholds. The detection rate varied across extraction methods, basecalling models, and primer choices, with BT consistently outperforming all other extraction workflows in terms of rapid species recovery (Figure 4). Statistical analysis confirms that BT significantly accelerated species accumulation, with full species recovery occurring within 3–5 hours across most workflows (GAM: edf = 3.908, χ² = 89.16, p < 2e-16). In contrast, Chelex required over 50 hours to detect the full species set, underscoring its inefficiency in DNA recovery (Figure 4). This delayed accumulation likely reflects lingering effects of PCR inhibitors that are not fully removed during cleanup, limiting downstream amplification despite post-PCR purification.

**Figure 4.**
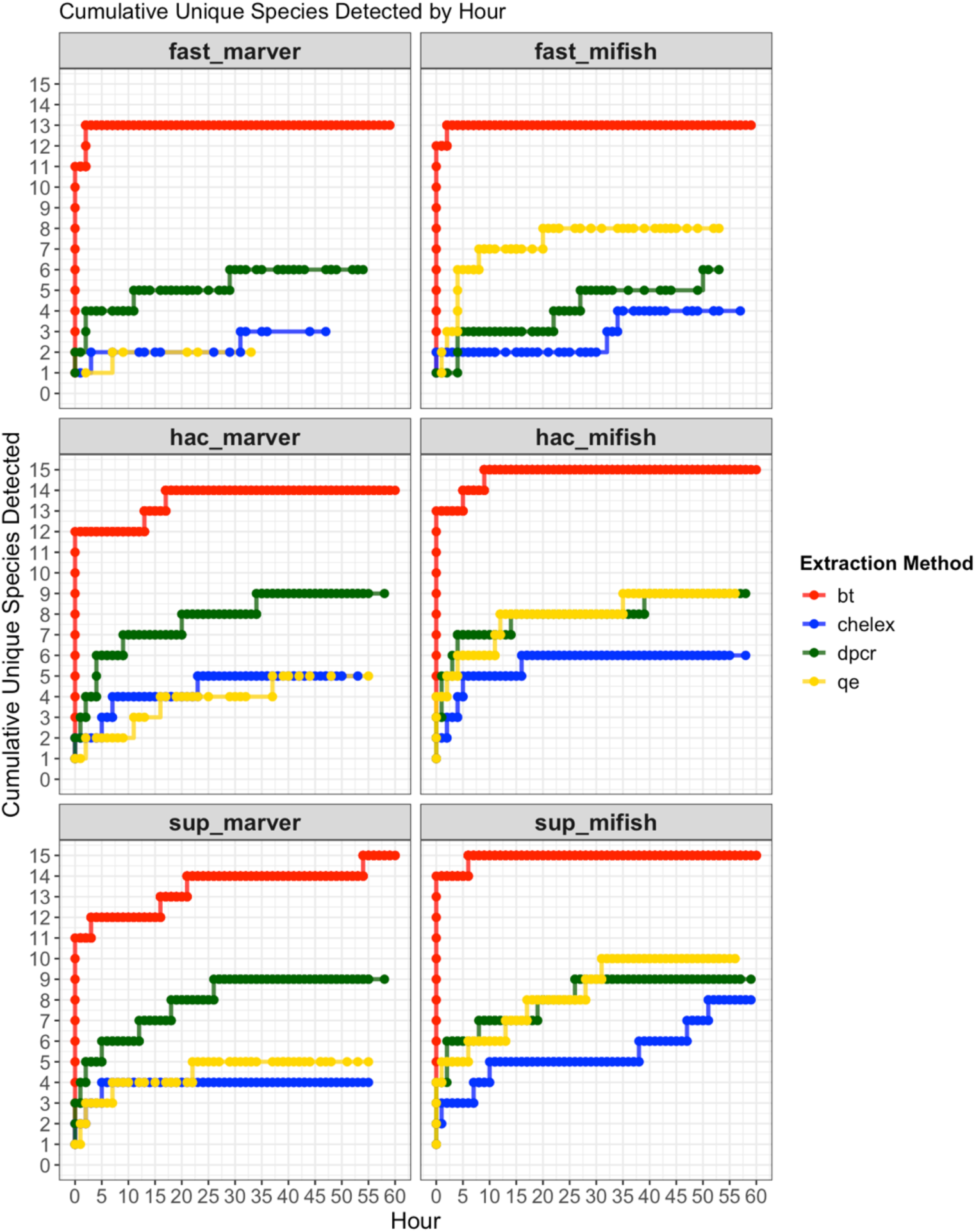
Cumulative Unique Species Detected Over Sequencing Time. The x-axis represents sequencing time in hours, while the y-axis indicates the cumulative number of detected species. Color indicates DNA extraction methods: Qiagen Blood & Tissue (BT, red), Chelex (blue), DirectPCR (dPCR, green), and QuickExtract (QE, yellow), demultiplexed using OBITools4. Each facet corresponds to a combination of basecalling model (FAST, HAC, SUP) and primer set (MiFish-U, MarVer1).

**Figure 5.**
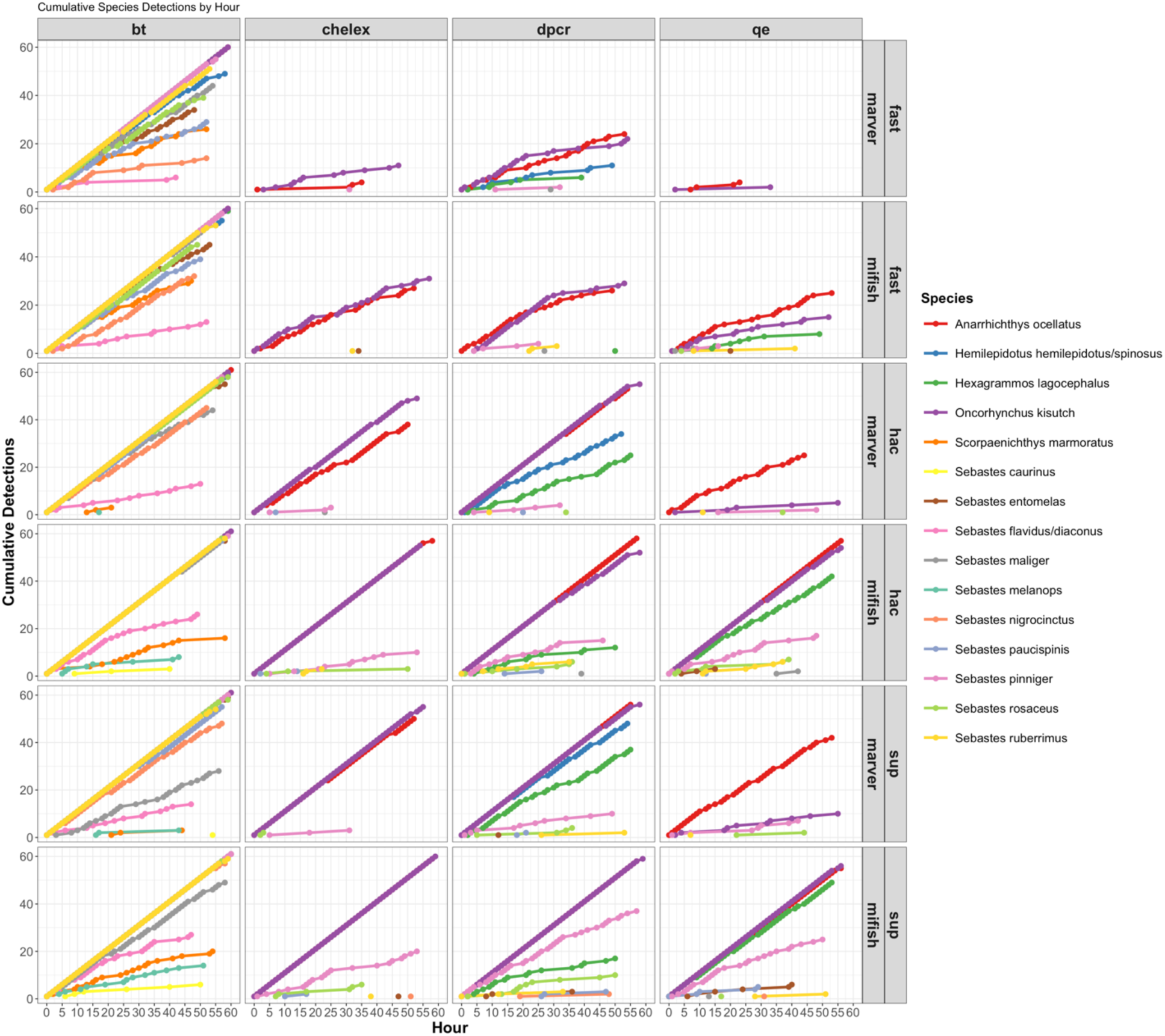
Cumulative Species Detections Over Sequencing Time. The x-axis represents sequencing time in hours, while the y-axis indicates the cumulative number of detections per species (color). The 24 panels represent variations in DNA extraction method (BT, Chelex, dPCR, QuickExtract), basecalling model (FAST, HAC, SUP), and primer set (MiFish-U, MarVer1).

BT-extracted samples, when paired with the MiFish-U primer set and SUP basecalling, reached a plateau at 15 species within 3 hours (Table 5), indicating optimal conditions for rapid species detection. Even under less efficient workflow conditions (FAST basecalling or MarVer1 primers), BT samples consistently outperformed other extraction methods, with ≥9 species detected within the first hour (Figure 4). The DirectPCR and QuickExtract extractions exhibited intermediate performance (near-complete detection within 10-12 hours), while Chelex-extracted samples detected only 6–7 species detected by the 24-hour mark and complete detection (>10 species) requiring over 50 hours. This trend is confirmed by ZOID posterior coefficients, which were consistently negative for Chelex across multiple species (Figure 3; Supplementary Figure S4).

**Table 5.** Time-to-detection across eDNA extraction methods.

| Extraction Method | Hours to Detect 15 Species | Hours to Detect 12 Species | Hours to Detect 9 Species |
| --- | --- | --- | --- |
| BT (Qiagen) | 3–5 hours | 1.5–3 hours | <1 hour |
| DirectPCR | 10–12 hours | 5–7 hours | 3 hours |
| QuickExtract | 10–12 hours | 5–7 hours | 3 hours |
| Chelex | >50 hours | ~24 hours | ~12 hours |

A more granular breakdown of cumulative species detections over sequencing time is demonstrated in Figure 5, with each line representing one of the 15 detected species. The accumulation patterns reveal clear differences between species detected early versus those that appeared later. Common species, such as *Oncorhynchus kisutch* (Coho salmon), *Sebastes caurinus* (Copper rockfish), and *Hexagrammos lagocephalus* (Rock Greenling), exhibited a near-linear accumulation curve across sequencing time, reflecting a 1:1 detection per hour rate in most workflows. These species were consistently detected early, often within the first 1–2 hours of sequencing, regardless of extraction method.

In contrast, several species, including *Sebastes ruberrimus* (Yelloweye rockfish), *Sebastes melanops* (Black rockfish), and *Scorpaenichthys marmoratus* (Cabezon), showed delayed and irregular detection patterns, with steep stepwise increments in their accumulation curves. These species often required >10 hours of sequencing to achieve stable detection, particularly in DirectPCR, QuickExtract, and Chelex workflows. The necessity for prolonged sequencing in these cases is attributed to their lower initial eDNA concentrations, requiring additional sequencing depth to detect residual molecules before depletion. While these trends suggest differences in eDNA abundance or amplification efficiency, we refer to taxa as “less common” solely based on their later or less consistent detection in the sequencing run, rather than confirmed template concentrations.

Interestingly, extended sequencing times (>40 hours) did not always contribute to increased species recovery in less efficient workflows (e.g., Chelex + FAST basecalling), as species detections began to plateau despite continued sequencing. This trend suggests that rare taxa may become depleted over time, a phenomenon observed in stochastic eDNA capture processes where fewer molecules remain available for sequencing as sampling progresses. This aligns with ONT’s single-molecule sequencing mechanism, where rare molecules—once sequenced—may become depleted, particularly in workflows with limited input DNA (Hook & Timp, 2023). The GAM model (edf = 3.908, χ² = 89.16, p < 2e-16) supports this interpretation, demonstrating a significant nonlinear relationship between sequencing time and species accumulation, with diminishing returns observed beyond 30–40 hours.

These results underscore the critical role of DNA extraction, sequencing depth, and basecalling models in optimizing eDNA workflows. The greater reliability of Qiagen BT workflows, demonstrated by their consistent and rapid species recovery, may offer added value in low-resource or high-throughput settings. In such contexts, minimizing time spent on troubleshooting and repeat runs is often more important than reducing per-sample costs. The data suggest that short-duration sequencing (3–6 hours) is sufficient for full species recovery when using BT extraction and MiFish-U primers, making it a robust choice for rapid eDNA applications. However, studies targeting rarer species may benefit from extended sequencing (10–12 hours), particularly in workflows using DirectPCR, QuickExtract, or MarVer1 primers. Beyond 40 hours, sequencing returns diminish significantly, with only marginal gains in species detections. This plateau effect is most pronounced in Chelex-extracted samples, where even prolonged sequencing fails to recover the full species set efficiently.

## 4. Discussion

### 4.1. Performance and Feasibility of Field-Adaptable Extraction Methods

Our findings reinforce the pivotal role of DNA extraction methods in eDNA workflows, influencing both species recovery and practical feasibility. Across all tested conditions, Qiagen BT outperformed other methods, consistently detecting up to 15 species in the controlled aquarium setting. We also demonstrated that BT showed the strongest positive posterior coefficients across species, confirming its reliability in maximizing species detection. This aligns with prior studies demonstrating that spin-column kits maximize DNA recovery across diverse conditions due to their silica-membrane purification, which efficiently binds DNA while removing PCR inhibitors (Kranzfelder et al., 2016; Majaneva et al., 2018).

In contrast, DirectPCR and QuickExtract, while capable of recovering ∼80% of species, lack targeted inhibitor removal, which may lead to reduced efficiency in high-turbidity or inhibitor-rich environments (Majaneva et al., 2018). Future refinements, such as additional inhibitor removal steps or pre-concentration techniques (e.g., QuickConc; Kuroita et al., 2024), could improve DNA yield in challenging conditions. QuickConc, for instance, uses benzalkonium chloride with dispersed glass fibers to concentrate DNA and has demonstrated promising yield improvements in early trials. Similarly, employing pre-filters or specialized extraction aids could enhance the usability of ultra-portable methods like QuickExtract, particularly in environments with high particulate loads (Majaneva et al., 2018; Lee-Rodriguez et al., 2024). Their ability to achieve comparable detection without the need for either cold-chain logistics or high-speed centrifugation makes them promising for field applications. However, DirectPCR still requires a small centrifuge, posing a barrier in ultra-remote settings lacking stable power or benchtop equipment (Ip et al., 2024; Kirtane & Deiner, 2024).

Among low-cost alternatives, Chelex was the least reliable, frequently missing rarer taxa, especially in inhibitor-rich conditions. This outcome is consistent with its non-selective binding properties, which lead to poor nucleic acid retention, and with known limitations of Chelex-based protocols, which often carry over PCR inhibitors and reduce amplification efficiency (Walsh et al., 1991; Karlsson et al., 2022; Bracken et al., 2019). Chelex is extremely cost-effective (∼$0.05 per sample) which makes it attractive for high-throughput applications, even though extraction kits such as BT and QuickExtract typically cost several-fold more. However, the trade-off between cost and sensitivity suggests the Chelex may be better suited to scenarios where only the detection of dominant taxa is sufficient.

Ultimately, no single extraction method is universally superior, as the choice depends on study objectives and resource constraints (Table 6). Further developments in passive filtration or power-free filtration systems (Bessey et al., 2021; Kuroita et al., 2024) could enhance eDNA workflow simplification in resource-limited settings.

**Table 6.**
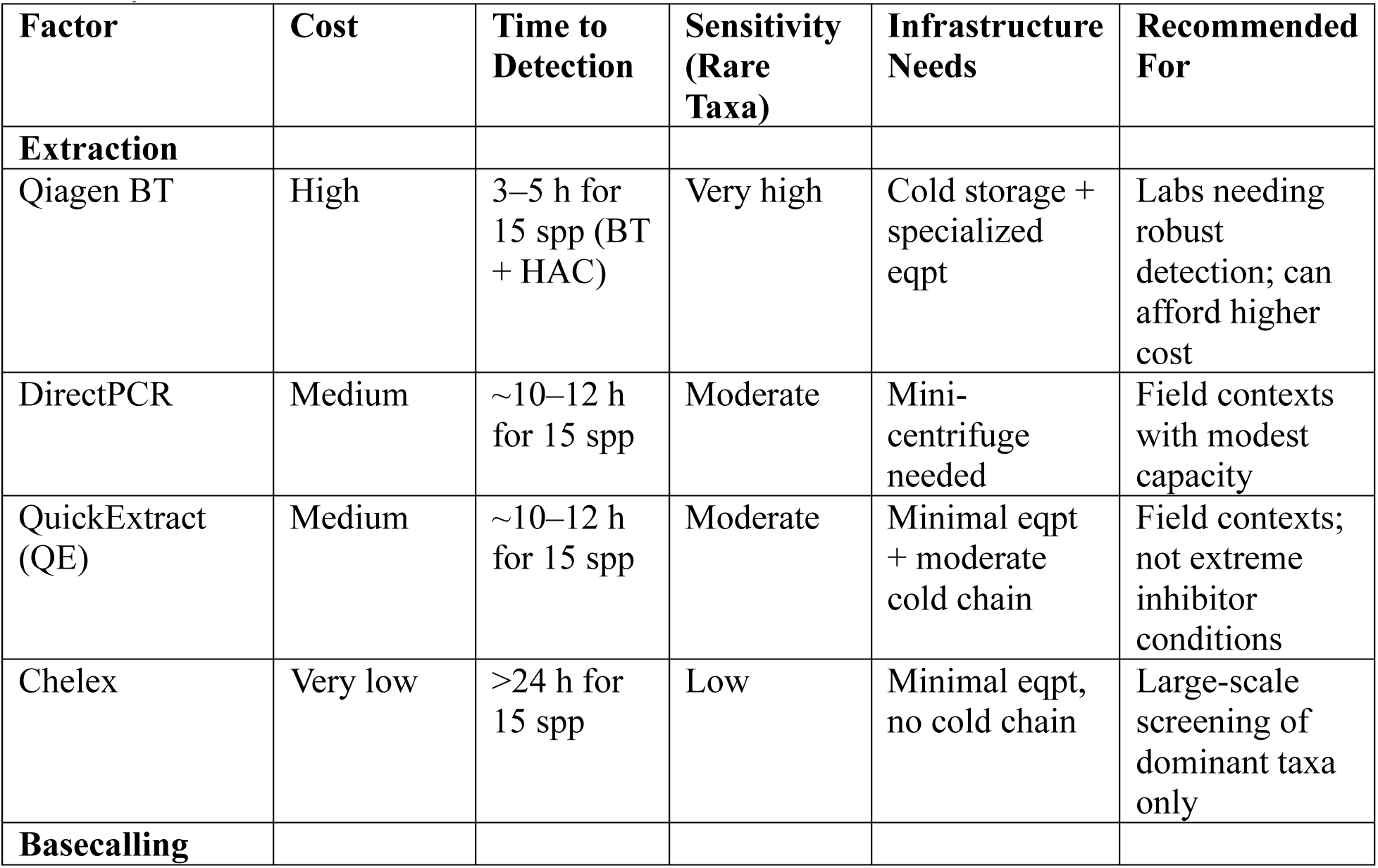
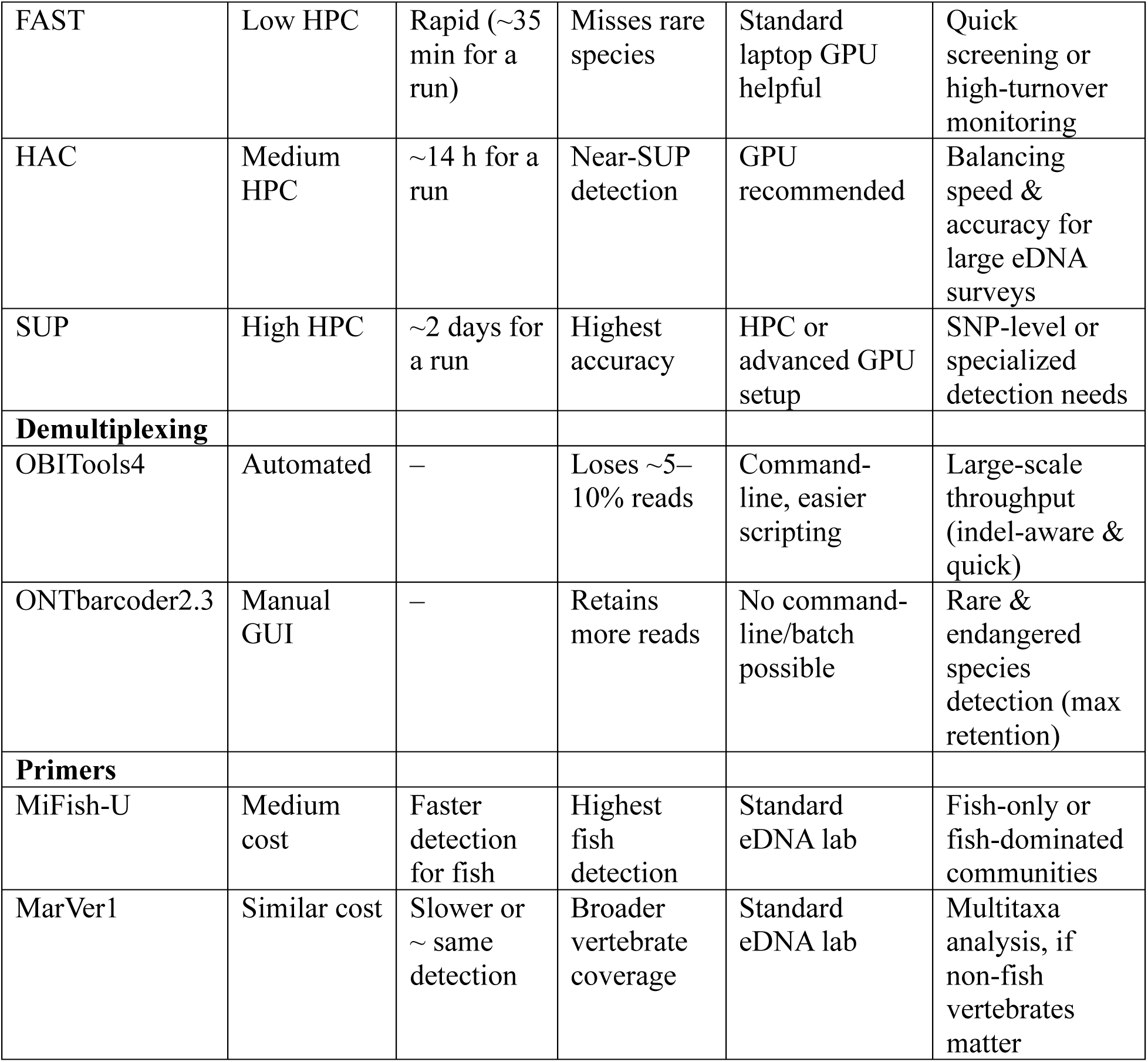
Decision Matrix for eDNA Workflow Optimization Based on Cost, Performance, and Suitability.

| Factor | Cost | Time to Detection | Sensitivity (Rare Taxa) | Infrastructure Needs | Recommended For |
| --- | --- | --- | --- | --- | --- |
| **Extraction** |  |  |  |  |  |
| Qiagen BT | High | 3–5 h for 15 spp (BT + HAC) | Very high | Cold storage + specialized eqpt | Labs needing robust detection; can afford higher cost |
| DirectPCR | Medium | ~10–12 h for 15 spp | Moderate | Mini-centrifuge needed | Field contexts with modest capacity |
| QuickExtract (QE) | Medium | ~10–12 h for 15 spp | Moderate | Minimal eqpt + moderate cold chain | Field contexts; not extreme inhibitor conditions |
| Chelex | Very low | >24 h for 15 spp | Low | Minimal eqpt, no cold chain | Large-scale screening of dominant taxa only |
| **Basecalling** |  |  |  |  |  |
| FAST | Low HPC | Rapid (~35 min for a run) | Misses rare species | Standard laptop GPU helpful | Quick screening or high-turnover monitoring |
| HAC | Medium HPC | ~14 h for a run | Near-SUP detection | GPU recommended | Balancing speed & accuracy for large eDNA surveys |
| SUP | High HPC | ~2 days for a run | Highest accuracy | HPC or advanced GPU setup | SNP-level or specialized detection needs |
| **Demultiplexing** |  |  |  |  |  |
| OBITools4 | Automated | – | Loses ~5–10% reads | Command-line, easier scripting | Large-scale throughput (indel-aware & quick) |
| ONTbarcode2.3 | Manual GUI | – | Retains more reads | No command-line/batch possible | Rare & endangered species detection (max retention) |
| **Primers** |  |  |  |  |  |
| MiFish-U | Medium cost | Faster detection for fish | Highest fish detection | Standard eDNA lab | Fish-only or fish-dominated communities |
| MarVer1 | Similar cost | Slower or ~ same detection | Broader vertebrate coverage | Standard eDNA lab | Multitaxa analysis, if non-fish vertebrates matter |

### 4.2. Trade-offs in Primer Selection: Specificity vs. Taxonomic Coverage

While DNA extraction determines the quantity and quality of eDNA recovered, primer selection further refines taxonomic resolution by influencing the breadth and specificity of species detection. As expected, the fish-specific MiFish-U primer set consistently outperformed MarVer1 in terms of read count, recovering approximately three times more fish reads per run. This aligns with previous studies demonstrating that primers explicitly designed for fish detection maximize on-target amplification and improve sequencing efficiency (Plewnia et al., 2024; Tibone et al., 2025).

MarVer1, despite yielding fewer total fish reads, detected most fish species and additionally recovered non-fish vertebrates, which may provide a more comprehensive view of community composition. In contrast, MiFish-U remains the optimal primer for high-confidence fish detection with minimal off-target amplification. In other words, the choice of primer depends on study objectives: for dedicated fish surveys, MiFish-U is preferred, whereas for ecosystem-wide studies where broader vertebrate diversity is important, MarVer1 can offer added value. Future research should explore whether combining multiple markers, such as 12S with mitochondrial D-loop regions (Andruszkiewicz et et al., 2020; Suarez-Bregua et al., 2022), can improve taxonomic resolution while preserving broad ecological coverage. Moreover, advancing long-read sequencing technologies may enable hybrid marker strategies that bridge the gap between specificity and inclusivity in species-level assignments. (Chang et al., 2024; Domingo-Bretón et al., 2024).

### 4.3. Computational Trade-Offs: Basecalling Accuracy and Demultiplexing Efficiency

Basecalling and demultiplexing are critical steps in eDNA workflows, each introducing trade-offs between speed, accuracy, and species recovery. Our results highlight how basecalling models impact detection efficiency, with FAST mode significantly reducing processing time (<1 hour) but at the cost of lower quality scores. These lower quality reads often lack the resolution needed for confident species-level identification, leading to higher uncertainty in taxonomic assignments and effectively dropped from downstream analyses, contributing to inconsistent detection of low-abundance taxa. HAC basecalling, in contrast, maintains a strong balance, recovering nearly all target species while completing basecalling within a reasonable timeframe (∼14 hours). Recent studies have demonstrated that HAC-optimized basecalling with R10.4.1 flow cells improve sequence accuracy by reducing homopolymer errors, which previously limited Nanopore species identification (Ni et al., 2023). Although SUP provides the highest read accuracy and retention, its extended processing time (two days to two weeks) makes it impractical for most field or real-time applications.

These findings align with previous research demonstrating that modern ONT basecallers, particularly when paired with R10.4.1 flow cells and updated chemistries, can approach Illumina-level accuracy (Chang et al., 2024; Stoeck et al., 2024; Dierickx et al., 2024). In rapid-response contexts, such as invasive species detection, FAST basecalling may be sufficient when only major taxa are of interest. However, for comprehensive biodiversity surveys targeting rare or cryptic species, HAC remains the preferred option, offering an optimal balance between speed and sensitivity. SUP may be necessary for specialized applications such as population genetics, where the highest basecalling fidelity is required.

Beyond basecalling, demultiplexing approaches further shape species recovery patterns, influencing whether low-abundance taxa are retained or lost. OBITools4, which trims primers, UMIs, and barcodes in a single command, provides a highly automated workflow suited for large-scale studies. While OBITools4 is indel aware and offers greater automation, this efficiency comes with trade-offs: consistently retaining fewer reads than ONTbarcoder2.3, potentially leading to missed detections of rare taxa. The observed difference in read retention (∼5–10% more for ONTbarcoder2.3) could directly impact whether a species is detected, particularly in eDNA samples with extremely low DNA concentrations.

For large-scale monitoring programs or studies requiring rapid processing of high-throughput sequencing data, OBITools4 remains a valuable tool for streamlining analysis. However, in scenarios where every read is critical, such as pathogen surveillance or the detection of endangered species, ONTbarcoder2.3’s more manual but higher-recovery approach may be preferable. Future refinements, such as adjusting OBITools4’s mismatch thresholds or integrating read-recovery scripts, could help bridge the gap between automation and sensitivity, ensuring that computational trade-offs do not compromise biodiversity assessments.

### 4.4. Detection Plateau in Real-Time eDNA Sequencing

Our hourly analysis over a 61-hour nanopore sequencing run revealed how quickly detection of the 15 known aquarium species plateaued under different workflows. In optimal scenarios (BT + MiFish-U + HAC), nearly all species were consistently detected within 3–5 hours. These short timeframes are particularly valuable for real-time ecological monitoring, as field teams could stop sequencing once a detection plateau is reached, conserving resources without compromising species recovery.

Conversely, suboptimal workflows showed a much slower species detection, often requiring ≥24 hours for detection to stabilize and >50 hours to recover only a partial species set. This highlights a direct link between method efficiency and sequencing duration, where lower DNA yields or increased sequencing errors can prolong the time needed to achieve full species detection. Extending sequencing time does not compensate for poor extraction efficiency. If DNA yields are too low or degraded at the start, additional sequencing contributes minimal new detections and only increases computational costs.

Our findings suggest that implementing adaptative stopping during sequencing protocols can optimize run times. For routine biodiversity surveys, sequencing 3–5 hours are sufficient using high-yield workflows (BT + MiFish-U + HAC). For low-abundance species, sequencing ∼10–12 hours may improve detection. Beyond 40 hours, further sequencing is unlikely to be cost-effective, as additional species detections plateau at <5% new recoveries. This plateau likely reflects a combination of library complexity, sequencing redundancy, and depletion of new informative DNA molecules, rather than a strict technological limitation. However, it is important to note that our study focused on detecting 15 target species. In broader biodiversity surveys, such as community-wide freshwater or marine assessments, extended sequencing might still yield new detections beyond 40 hours, particularly for rare or low-abundance taxa. These insights reinforce the idea that real-time sequencing analytics can optimize run duration dynamically, allowing researchers to adaptively terminate sequencing once species accumulation stabilizes, thereby reducing costs and computational burden (Chang et al., 2020; Plewnia et al., 2024).

### 4.5. Toward Accessible and Scalable eDNA Monitoring in Resource-Limited Settings

Given these findings, we outline key considerations for scaling eDNA workflows to resource-limited settings, focusing on logistics, and methodological accessibility. Our data show that simplified extractions (DirectPCR, QuickExtract) and high accuracy basecalling (HAC) can still recover most known taxa, making them viable alternatives when working under budget or infrastructure constraints. Notably, BT workflows may also offer added value in low-resource settings by minimizing troubleshooting and reruns, which can outweigh per-sample cost savings. These results align with previous studies demonstrating that Nanopore-based metabarcoding can approximate Illumina’s performance at lower cost and with reduced laboratory requirements (Tibone et al., 2025; Veillat et al., 2024; Stoeck et al., 2024; Kirchgeorg et al., 2024).

From a capacity-building perspective, integrating cost-effective extractions (DirectPCR, QuickExtract) with portable MinION sequencers in a “lab-in-a-box” format (Watsa et al., 2020; Chang et al., 2020) could significantly expand global eDNA adoption, particularly in biodiversity monitoring programs in the Global South, where infrastructure and technical expertise may be limited (Hirsch et al., 2024). In addition to logistical benefits, localized sequencing enables greater data sovereignty, allowing researchers to control their own workflows and analyses without relying on centralized sequencing facilities. Recent hardware advancements, such as The Oxford Nanopore MinION Mk1D, combined with its low cost, further enhance the feasibility of field-based sequencing by allowing direct sequencing on an iPad, eliminating the need for a high-performance laptop and making remote sequencing even more accessible. In field settings, fast basecalling is the most practical option for real-time applications, as more intensive models may generate analysis lag due to computational constraints on less powerful computing devices.

Despite these advances, several challenges remain in making eDNA workflows fully field deployable. Achieving cold-chain independence, standardizing protocols for inhibitor-rich samples, and ensuring adequate training on simplified extraction methods are ongoing barriers (Rieder et al., 2024). While ONT sequencing can be performed offline using preconfigured settings, internet access is still required for license verification, software updates, and troubleshooting, limiting the feasibility of fully independent field deployments. Expanding support for offline configurations would further enhance the portability of eDNA sequencing (Geckeler et al., 2025 in review). Addressing these constraints through clear standard operating procedures (SOPs), remote training programs, and improved sample preservation techniques will be critical for broader implementation (Hirsch et al., 2024; Rieder et al., 2024; Stammnitz et al., 2024).

Beyond sequencing, bioinformatics accessibility remains a major hurdle for eDNA-users without coding training. Many analysis pipelines rely on command-line tools, limiting adoption in non-specialized labs. However, cloud-based platforms, graphical user interface (GUI)-driven tools and low-cost field-deployable computational resources are emerging as viable alternatives (Bloomfield et al., 2024; Ip et al., 2023). Recent advancements in affordable, low-power GPUs and edge computing devices now enable real-time basecalling and analysis on a budget. Bloomfield et al. (2024) demonstrated that Nanopore sequencing could be implemented without on-site bioinformaticians using a GPU costing only $649, highlighting the feasibility of decentralized, cost-effective sequencing analysis. Such approaches, when combined with cloud-based Galaxy-tools bioinformatics platforms (Martin et al., 2024) and workflow automation software like APSCALE (Buchner et al., 2022), significantly lower financial barriers, computational expertise and hardware barriers for non-experts. Similarly, CrocoBLAST (Boratyn et al., 2017) provides an online GUI classifier, enabling species identification without the need for command-line Kraken2, BLASTn, or other taxonomic assignment tools. These advancements are instrumental in bridging the bioinformatics accessibility gap, making low-cost, scalable eDNA analysis more inclusive. Our study directly evaluates these simplified approaches, providing empirical support for their viability in resource-limited field settings.

Emerging low-power, field-adaptable molecular tools could further enhance eDNA accessibility. Isothermal amplification methods (e.g., RPA) offer an attractive alternative to traditional PCR, reducing power demands and simplifying workflow requirements (Plewnia et al., 2025). When integrated with in situ sequencing, such innovations could enable near real-time eDNA assessments, expanding the feasibility of biodiversity monitoring in remote environments. While such approaches were beyond the scope of this study, they represent logical next steps in the continued democratization of eDNA methodologies, ensuring that species monitoring remains scalable, cost-effective, and accessible, even in the most infrastructure-limited environments.

## 5. Conclusion

By evaluating 144 workflow combinations in a controlled aquarium system, this study provides a comprehensive assessment of methodological trade-offs in eDNA monitoring. Rather than prescribing a single “best” approach, our findings highlight key decision points where optimizations can improve efficiency without disproportionately sacrificing species recovery. Among extraction methods, BT remains the most reliable, detecting ∼15 species within 5 hours, while DirectPCR and QuickExtract offer field-adaptable alternatives that require 10–12 hours but function with minimal infrastructure. Chelex, though cost-effective, exhibits reduced sensitivity and requires >24 hours to reach close-to-similar detection levels. Computationally, HAC basecalling balances speed and accuracy, and MiFish-U enhances fish-specific detection, while MarVer1 broadens taxonomic scope, making it better suited for multi-taxa assessments. Demultiplexing further affects read retention, with ONTbarcoder2.3 preserving more low abundance reads and OBITools4 offering automation at the cost of slightly lower sensitivity. Real-time sequencing analysis suggests that most optimized workflows plateau in 3–5 hours, supporting the potential for adaptive stopping strategies to reduce sequencing costs without compromising detection. Our Bayesian zero-and-one-inflated Dirichlet model explicitly accounts for variation at both the PCR-replicate and biological-replicate levels, and confirms that the combination of BT extraction, MiFish-U primers, and ONTbarcoder2.3 demultiplexing yield the highest detection odds across species. However, both DirectPCR and QuickExtract also performed reasonably across most species, offering competitive detection rates in workflows with minimal lab infrastructure.

Overall, these results illustrate that no single workflow is universally optimal, but rather that each method offers distinct advantages that can be matched to specific study goals. These findings provide a transparent decision framework for researchers to navigate trade-offs between detection sensitivity, cost efficiency, and sequencing speed based on study constraints. By identifying where meaningful losses occur, we highlight the conditions under which cost-effective or rapid workflows can maintain reliable species detection and where they risk compromising accuracy. Rather than simply making eDNA workflows cheaper or faster, our goal was to clarify what is gained or lost with each methodological choice, ensuring that researchers can optimize their workflows based on project priorities. By clearly mapping the trade-offs between cost, time, and detection accuracy, this study empowers researchers to make informed methodological choices, ensuring that eDNA monitoring remains scalable, efficient, and adaptable across diverse ecological and logistical settings.

## 6. Acknowledgments

We thank the Seattle Aquarium for assisting with the water sampling and for their logistical support. We also acknowledge OceanKind [Grant No. GR042390] and Packard Foundation [Grant No. GR016745] for funding support and the Centre of Environmental Genomics for access to HPC resources. Special thanks to Olivia M. Scott for the field work assistance, Megan R. Shaffer for early discussions on PCR optimizations, and Ole Shelton for the data visualization advice.

## Author Contributions

- Conceptualization: Y.C.A.I, E.A.A, R.P.K
- Methodology: Y.C.A.I, E.A.A.
- Formal Analysis: Y.C.A.I, with inputs from E.A.A., R.P.K
- Writing—Original Draft: Y.C.A.I.
- Writing—Review & Editing: E.A.A, R.P.K, S.L.H.
- Funding Acquisition: E.A.A, R.P.K

## Data Availability

Raw FASTQ files have been deposited in the NCBI Sequence Read Archive under BioProject XXX. All other data supporting the findings of this study are provided in the main text and its supplementary materials.

## Conflict of Interest

The authors declare no conflict of interest.

## Notes

### Competing Interest Statement

The authors have declared no competing interest.

## 7. References

Allan, E. A., Kelly, R. P., D’Agnese, E. R., Garber-Yonts, M. N., Shaffer, M. R., Gold, Z. J., & Shelton, A. O. (2023). Quantifying impacts of an environmental intervention using environmental DNA. Ecological Applications, 33(8), e2914.

Andruszkiewicz, E. A., Yamahara, K. M., Closek, C. J., & Boehm, A. B. (2020). Quantitative PCR assays to detect whales, rockfish, and common murre environmental DNA in marine water samples of the Northeastern Pacific. PloS One, 15(12), e0242689.

Bayer, P. E., Bennett, A., Nester, G., Corrigan, S., Raes, E. J., McInnes, A. S.,… & Rauschert, S. (2024). A comprehensive evaluation of taxonomic classifiers in marine vertebrate eDNA studies. bioRxiv, 2024–02.

Bloomfield, M., Hutton, S., Velasco, C., Burton, M., Benton, M., & Storey, M. (2024). Oxford Nanopore next generation sequencing in a front-line clinical microbiology laboratory without on-site bioinformaticians. Pathology, 56(3), 444–447.

Boyer, F., Mercier, C., Bonin, A., Le Bras, Y., Taberlet, P., & Coissac, E. (2016). obitools: A unix-inspired software package for DNA metabarcoding. Molecular ecology resources, 16(1), 176–182.

Bracken, F. S., Rooney, S. M., Kelly-Quinn, M., King, J. J., & Carlsson, J. (2019). Identifying spawning sites and other critical habitat in lotic systems using eDNA “snapshots”: A case study using the sea lamprey *Petromyzon marinus* L. Ecology and Evolution, 9(1), 553–567.

Buchner, D., Macher, T. H., & Leese, F. (2022). APSCALE: advanced pipeline for simple yet comprehensive analyses of DNA metabarcoding data. Bioinformatics, 38(20), 4817–4819.

Chang, J. J. M., Ip, Y. C. A., Neo, W. L., Mowe, M. A., Jaafar, Z., & Huang, D. (2024). Primed and ready: nanopore metabarcoding can now recover highly accurate consensus barcodes that are generally indel-free. BMC Genomics, 25(1), 842.

Chang, J. J. M., Ip, Y. C. A., Ng, C. S. L., & Huang, D. (2020). Takeaways from mobile DNA barcoding with BentoLab and MinION. Genes, 11(10), 1121.

Deiner, K., Bik, H. M., Mächler, E., Seymour, M., Lacoursière-Roussel, A., Altermatt, F.,… & Bernatchez, L. (2017). Environmental DNA metabarcoding: Transforming how we survey animal and plant communities. Molecular Ecology, 26(21), 5872–5895.

Dierickx, G., Tondeleir, L., Asselman, P., Vandekerkhove, K., & Verbeken, A. (2024). What Quality Suffices for Nanopore Metabarcoding? Reconsidering Methodology and Ectomycorrhizae in Decaying *Fagus sylvatica* Bark as Case Study. Journal of Fungi, 10(10), 708.

Domingo-Bretón, R., Moroni, F., Toxqui-Rodríguez, S., Belenguer, Á., Piazzon, M. C., Pérez-Sánchez, J., & Naya-Català, F. (2024). Moving Beyond Oxford Nanopore Standard Procedures: New Insights from Water and Multiple Fish Microbiomes. International Journal of Molecular Sciences, 25(23), 12603.

Doorenspleet, K., Jansen, L., Oosterbroek, S., Kamermans, P., Bos, O., Wurz, E.,… & Nijland, R. (2025). The Long and the Short of It: Nanopore-Based eDNA Metabarcoding of Marine Vertebrates Works; Sensitivity and Species-Level Assignment Depend on Amplicon Lengths. Molecular Ecology Resources, e14079.

Ficetola, G. F., & Taberlet, P. (2023). Towards exhaustive community ecology via DNA metabarcoding. Molecular Ecology, 32(23), 6320–6329.

Geckeler, C., Kirchgeorg, S., Strunck, G., Thostrup, F. B., Sangermano, F., Desiderato, A., Lüthi, M., Jucker, M., Gonzalez Herrera, M. A., Franco Sierra, N. D., Pulido-Santacruz, P., Jin Marc, C. J., Ip, Y. C. A., Mä chler, E., Svenning, A., Mougeot, G., Høye, T., Fopp, F., Pellissier, L., Dao, D., Deiner, K., Melvad, C., Hamaza, S., & Mintchev, S. (2025 in review). Field deployment of BiodivX drones in the Amazon rainforest for biodiversity monitoring. IEEE Journal of Field Robotics, xx(xx), xx–xx. xxxxx

Hirsch, S., Acharya-Patel, N., Amamoo, P. A., Borrero-Pérez, G. H., Cahyani, N. K. D., Ginigini, J. G.,… & Kelly, R. (2024). Centering accessibility, increasing capacity, and fostering innovation in the development of international eDNA standards. Metabarcoding and Metagenomics, 8, e126058.

Holman, L. E., Hollenbeck, C. M., Ashton, T. J., & Johnston, I. A. (2019). Demonstration of the use of environmental DNA for the non-invasive genotyping of a bivalve mollusk, the European flat oyster (*Ostrea edulis*). Frontiers in Genetics, 10, 1159.

Hook, P. W., & Timp, W. (2023). Beyond assembly: the increasing flexibility of single-molecule sequencing technology. Nature Reviews Genetics, 24(9), 627–641.

Ip, Y. C. A., Chang, J. J. M., & Huang, D. (2023). Chapter Advancing and Integrating ‘Biomonitoring 2.0’with New Molecular Tools for Marine Biodivesity and Ecosystem Assessments. Oceanography and Marine Biology.

Ip, Y. C. A., Chang, J. J. M., Lim, K. K., Jaafar, Z., Wainwright, B. J., & Huang, D. (2021). Seeing through sedimented waters: environmental DNA reduces the phantom diversity of sharks and rays in turbid marine habitats. BMC Ecology and Evolution, 21, 1–14.

Ip, Y. C. A., Chen, J., Tan, L. Y., Lau, C., Chan, Y. H., Shanmugavelu Balasubramaniam, R.,… & Yap, H. H. (2024). Establishing environmental DNA and RNA protocols for the simultaneous detection of fish viruses from seawater. Environmental DNA, 6(1), e418.

Jensen, A. J., Kelly, R. P., Anderson, E. C., Satterthwaite, W. H., Shelton, A. O., & Ward, E. J. (2022). Introducing zoid: A mixture model and R package for modeling proportional data with zeros and ones in ecology. Ecology, 103(11), e3804.

Karlsson, E., Ogonowski, M., Sundblad, G., Sundin, J., Svensson, O., Nousiainen, I., & Vasemägi, A. (2022). Strong positive relationships between eDNA concentrations and biomass in juvenile and adult pike (*Esox lucius*) under controlled conditions: Implications for monitoring. Environmental DNA, 4(4), 881–893.

Kirchgeorg, S., Chang, J. J. M., Ip, Y. C. A., Jucker, M., Geckeler, C., Lüthi, M.,… & Mintchev, S. (2024). eProbe: sampling of environmental DNA within tree canopies with drones. Environmental Science & Technology, 58(37), 16410–16420.

Kirtane, A., & Deiner, K. (2024). Improving the recovery for dissolved eDNA state: A comparative analysis of isopropanol precipitation, magnetic bead extraction, and centrifugal dialysis. Environmental DNA, 6(3), e572.

Kranzfelder, P., Ekrem, T., & Stur, E. (2016). Trace DNA from insect skins: a comparison of five extraction protocols and direct PCR on chironomid pupal exuviae. Molecular Ecology Resources, 16(1), 353–363.

Krehenwinkel, H., Pomerantz, A., & Prost, S. (2019). Genetic biomonitoring and biodiversity assessment using portable sequencing technologies: current uses and future directions. Genes, 10(11), 858.

Kuroita, T., Iwamoto, R., Wu, Q., & Minamoto, T. (2024). QuickConc: A rapid, efficient, and power-free eDNA concentration method with cationic-assisted capture. Preprint.

Laamanen, T., Norros, V., Vihervaara, P., Jerney, J., Kortelainen, P., Kujala, K.,… & Meissner, K. (2025). Technology Readiness Level of biodiversity monitoring with molecular methods–where are we on the road to routine implementation? Metabarcoding and Metagenomics, 9, e130834.

Lee-Rodriguez, J., Ranger, C. M., Leach, A., Michel, A., Reding, M. E., & Canas, L. (2024). Using Environmental DNA to Detect and Identify Sweetpotato Whitefly *Bemisia argentifolii* and Twospotted Spider Mite *Tetranychus urticae* in Greenhouse-Grown Tomato Plants. Environmental DNA, 6(5), e70026.

Lepuschitz, S., Weinmaier, T., Mrazek, K., Beisken, S., Weinberger, J., & Posch, A. E. (2020). Analytical performance validation of next-generation sequencing based clinical microbiology assays using a K-mer analysis workflow. Frontiers in Microbiology, 11, 1883.

Liu, Y., Ghaffari, M. H., Ma, T., & Tu, Y. (2024). Impact of database choice and confidence score on the performance of taxonomic classification using Kraken2. aBIOTECH, 1–11.

Lu, J., Rincon, N., Wood, D. E., Breitwieser, F. P., Pockrandt, C., Langmead, B.,… & Steinegger, M. (2022). Metagenome analysis using the Kraken software suite. Nature Protocols, 17(12), 2815–2839.

Majaneva, M., Diserud, O. H., Eagle, S. H., Hajibabaei, M., & Ekrem, T. (2018). Choice of DNA extraction method affects DNA metabarcoding of unsorted invertebrate bulk samples. Metabarcoding and Metagenomics, 2, e26664.

McElreath, R. (2018). Statistical rethinking: A Bayesian course with examples in R and Stan. Chapman and Hall/CRC.

Miya, M., Sato, Y., Fukunaga, T., Sado, T., Poulsen, J. Y., Sato, K.,… & Iwasaki, W. (2015). MiFish, a set of universal PCR primers for metabarcoding environmental DNA from fishes: detection of more than 230 subtropical marine species. Royal Society open science, 2(7), 150088.

Ni, Y., Liu, X., Simeneh, Z. M., Yang, M., & Li, R. (2023). Benchmarking of Nanopore R10. 4 and R9. 4.1 flow cells in single-cell whole-genome amplification and whole-genome shotgun sequencing. Computational and Structural Biotechnology Journal, 21, 2352–2364.

O’Donnell, J. L., Kelly, R. P., Lowell, N. C., & Port, J. A. (2016). Indexed PCR primers induce template-specific bias in large-scale DNA sequencing studies. PloS One, 11(3), e0148698.

Petit-Marty, N., Casas, L., & Saborido-Rey, F. (2023). State-of-the-art of data analyses in environmental DNA approaches towards its applicability to sustainable fisheries management. Frontiers in Marine Science, 10, 1061530.

Plewnia, A., Krehenwinkel, H., & Heine, C. (2024). Towards low-cost and PCR free field-based community metabarcoding. Research Square Preprint.

Rieder, J., Jemmi, E., Hunter, M. E., & Adrian-Kalchhauser, I. (2024). A Guide to Environmental DNA Extractions for Non-Molecular Trained Biologists, Ecologists, and Conservation Scientists. Environmental DNA, 6(5), e70002.

Scriver, M., von Ammon, U., Youngbull, C., Pochon, X., Stanton, J. A. L., Gemmell, N. J., & Zaiko, A. (2024). Drop it all: extraction-free detection of targeted marine species through optimized direct droplet digital PCR. PeerJ, 12, e16969.

Shaffer, M. R., Allan, E. A., Van Cise, A. M., Parsons, K. M., Shleton, A. O., Kelly, R. P (2025, in press) Observation bias in metabarcoding. Molecular Ecology Resources.

Srivathsan, A., Feng, V., Suárez, D., Emerson, B., & Meier, R. (2024). ONTbarcoder 2.0: rapid species discovery and identification with real-time barcoding facilitated by Oxford Nanopore R10. 4. Cladistics, 40(2), 192–203.

Stammnitz, M. R., Hartman Scholz, A., & Duffy, D. J. (2024). Environmental DNA without borders: Let’s embrace decentralised genomics to meet the UN’s biodiversity targets. EMBO Reports, 25(10), 4095–4099.

Stoeck, T., Katzenmeier, S. N., Breiner, H. W., & Rubel, V. (2024). Nanopore duplex sequencing as an alternative to Illumina MiSeq sequencing for eDNA-based biomonitoring of coastal aquaculture impacts. Metabarcoding and Metagenomics, 8, e121817.

Suarez-Bregua, P., Álvarez-González, M., Parsons, K. M., Rotllant, J., Pierce, G. J., & Saavedra, C. (2022). Environmental DNA (eDNA) for monitoring marine mammals: Challenges and opportunities. Frontiers in Marine Science, 9, 987774.

Tibone, M., Stefanni, S., Aguzzi, J., O’Neill, B., & Mirimin, L. (2025). A multi-marker fish eDNA metabarcoding study comparing Illumina and Nanopore high-throughput sequencing platforms. Authorea Preprint.

Valsecchi, E., Bylemans, J., Goodman, S. J., Lombardi, R., Carr, I., Castellano, L.,… & Galli, P. (2020). Novel universal primers for metabarcoding environmental DNA surveys of marine mammals and other marine vertebrates. Environmental DNA, 2(4), 460–476.

Veillat, L., Boyer, S., Querejeta, M., Magnoux, E., Roques, A., Lopez-Vaamonde, C., & Roux, G. (2024). Benchmarking three DNA metabarcoding technologies for efficient detection of non-native cerambycid beetles in trapping collections. NeoBiota, 96, 237–259.

Walsh, P. S., Metzger, D. A., & Higuchi, R. (1991). Chelex 100 as a medium for simple extraction of DNA for PCR-based typing from forensic material. Biotechniques, 10(4), 506–513.

Wang, Z., Liu, X., Liang, D., Wang, Q., Zhang, L., & Zhang, P. (2023). VertU: universal multilocus primer sets for eDNA metabarcoding of vertebrate diversity, evaluated by both artificial and natural cases. Frontiers in Ecology and Evolution, 11, 1164206.

Watsa, M., Erkenswick, G. A., Pomerantz, A., & Prost, S. (2020). Portable sequencing as a teaching tool in conservation and biodiversity research. PLoS Biology, 18(4), e3000667.

Wick, R. R., Judd, L. M., & Holt, K. E. (2019). Performance of neural network basecalling tools for Oxford Nanopore sequencing. Genome Biology, 20, 1–10.

Wickham, H., Averick, M., Bryan, J., Chang, W., McGowan, L. D. A., François, R.,… & Yutani, H. (2019). Welcome to the Tidyverse. Journal of Open Source Software, 4(43), 1686.

Wood, S. N. (2001). mgcv: GAMs and generalized ridge regression for R. R news, 1(2), 20–25.

Wood, D. E., Lu, J., & Langmead, B. (2019). Improved metagenomic analysis with Kraken 2. Genome Biology, 20, 1–13.

Yang, J., Li, C., Lo, L. S. H., Zhang, X., Chen, Z., Gao, J.,… & Cheng, J. (2024). Artificial Intelligence-Assisted Environmental DNA Metabarcoding and High-Resolution Underwater Optical Imaging for Noninvasive and Innovative Marine Environmental Monitoring. Journal of Marine Science and Engineering, 12(10), 1729.

Zhu, T., Sato, Y., Sado, T., Miya, M., & Iwasaki, W. (2023). MitoFish, MitoAnnotator, and MiFish Pipeline: updates in 10 years. Molecular Biology and Evolution, 40(3), msad035.

